# Convergence of autism proteins at the cilium

**DOI:** 10.1101/2024.12.05.626924

**Authors:** Elina Kostyanovskaya, Micaela C. Lasser, Belinda Wang, James Schmidt, Ethel Bader, Chad Buteo, Juan Arbelaez, Aria Rani Sindledecker, Kate E. McCluskey, Octavio Castillo, Sheng Wang, Jeanselle Dea, Kathryn A. Helde, J. Michael Graglia, Elise Brimble, David B. Kastner, Aliza T. Ehrlich, Matthew W. State, A. Jeremy Willsey, Helen Rankin Willsey

**Affiliations:** Department of Psychiatry and Behavioral Sciences, UCSF Weill Institute for Neurosciences, University of California San Francisco, San Francisco, CA; SynGap Research Fund, Mill Valley, CA; Citizen Health, San Francisco, CA; Chan Zuckerberg Biohub – San Francisco, San Francisco, CA

## Abstract

Hundreds of high-confidence autism genes have been identified, yet the relevant etiological mechanisms remain unclear. Gene ontology analyses have repeatedly identified enrichment of proteins with annotated functions in gene expression regulation and neuronal communication. However, proteins are often pleiotropic and these annotations are inherently incomplete. Our recent autism functional genetics work has suggested that these genes may share a common mechanism at the cilium, a membrane-bound organelle critical for neurogenesis, brain patterning, and neuronal activity–all processes strongly implicated in autism. Moreover, autism commonly co-occurs with conditions that are known to involve ciliary-related pathologies, including congenital heart disease, hydrocephalus, and blindness. However, the role of autism genes at the cilium has not been systematically investigated. Here we demonstrate that autism proteins spanning disparate functional annotations converge in expression, localization, and function at cilia, and that patients with pathogenic variants in these genes have cilia-related co-occurring conditions and biomarkers of disrupted ciliary function. This degree of convergence among genes spanning diverse functional annotations strongly suggests that cilia are relevant to autism, as well as to commonly co-occurring conditions, and that this organelle should be explored further for therapeutic potential.

**One-Sentence Summary:** Autism genes of diverse functional annotations converge at cilia, deepening our understanding of underlying autism biology and co-occurring conditions.

## Introduction

Despite substantial progress in identifying genetic contributors to autism, translating success in gene discovery to an actionable understanding of underlying etiological mechanisms has been challenging (Willsey et al. 2022; Sestan and State 2018; A. J. Willsey et al. 2018; Willsey and State 2015). Gene ontology analyses of autism-associated genes have repeatedly identified enrichment of proteins with annotated functions in two predominant categories: gene expression regulation and neuronal communication (De Rubeis et al. 2014; O’Roak et al. 2012; Gilman et al. 2011; Iossifov et al. 2014; Satterstrom et al. 2020; Sanders et al. 2015). However, proteins are often highly pleiotropic and brain gene expression is particularly dynamic developmentally; therefore identifying which function(s) is/are relevant to autism itself has been difficult (Sestan and State 2018; A. J. Willsey et al. 2018; State and Šestan 2012; Willsey and State 2015). Our recent functional genetics work has highlighted this issue: autism genes annotated in neuronal communication also function during neurogenesis (Sun et al. 2024; Willsey et al. 2021); autism genes annotated in RNA splicing also function on cilia (Willsey et al. 2020); and autism genes annotated as chromatin regulators also function at microtubules of the mitotic centrosome (Lasser et al. 2023). Despite these various pleiotropic functions, our recent *in vivo* work in *Xenopus* and *in vitro* work in human neural progenitors intersect to implicate microtubule biology, specifically at the cilium, for several autism genes of diverse annotated function (Sun et al. 2024; Willsey et al. 2020, 2018; Teerikorpi et al. 2024; Willsey et al. 2021). This has led us to hypothesize that a substantial subset of autism genes may converge at the cilium, a microtubule-based organelle critical for neurogenesis, brain patterning, and neuronal excitability, all processes strongly implicated in autism (Packer 2016; Willsey et al. 2022). When reliable neuropsychiatric gene discovery was in its infancy, *in vitro* work implicated several genes, that are now considered high-confidence autism genes, in ciliation (Marley and von Zastrow 2012), but this possibility has not subsequently been explored systematically.

Cilia are membrane-bound organelles present on almost all cells, including neurons, with essential roles in intercellular communication, excitability, and signaling (Reiter and Leroux 2017; Hilgendorf et al. 2024). They are microtubule-based structures, nucleated by a modified centrosome, termed the basal body, with surrounding pericentriolar material and centriolar satellites (Odabasi et al. 2020; Bärenz et al. 2011; Kumar and Reiter 2021). Cilia are rich in GPCRs, ion channels, and signaling molecules, much like neuronal synapses, and similarly actively transport these proteins both within the cilium and also into the cilium across a diffusion barrier distal to the basal body (Park and Leroux 2022; Nechipurenko et al. 2013; Hilgendorf et al. 2024). Cilia are generally classified as either non-motile primary cilia that signal (as in neurons) or motile cilia that generate fluid flow (as in respiratory tract epithelial cells). In neurons, acute cell-autonomous loss of ciliary signaling or cilia results in a disruption of the balance between excitatory and inhibitory transmission by regulating excitatory synapses (Tereshko et al 2021). Ciliary dysfunction causes a wide spectrum of multi-organ clinical conditions including congenital heart defects, chronic respiratory issues, hypotonia, hydrocephalus, blindness, hearing loss, epilepsy, renal anomalies, dysmorphic features, and skeletal abnormalities (Gabriel et al. 2018; Djenoune et al. 2022; Klena et al. 2017; Reiter and Leroux 2017).

Consistently, autism commonly co-occurs with these cilia-related disorders including congenital heart disease (CHD), epilepsy, hydrocephalus, and blindness (Sigmon et al. 2019; Rosenthal et al. 2021; Al-Beltagi 2021; A. J. Willsey et al. 2018; Jin et al. 2017; Pili et al. 2021; Duy, Greenberg, et al. 2022; Homsy et al. 2015) and orthogonal gene discovery efforts, focused on highly penetrant rare mutations, have identified high-confidence autism genes among the risk genes associated with many of these conditions—particularly epilepsy and CHD (Homsy et al. 2015; Zaidi et al. 2013; Jin et al. 2017; A. J. Willsey et al. 2018; Rosenthal et al. 2021; Epi25 Collaborative 2024). Similarly, individuals with genetic ciliopathies such as Bardet-Biedl Syndrome and Joubert Syndrome (caused by variants in cilia-specific genes) have been reported to have a high prevalence of autistic features (Kerr et al. 2016; Ozonoff et al. 1999), supporting the possibility that cilia underlie a subset of autism-relevant biology. Based on these converging lines of evidence, we hypothesized that ciliary biology is relevant to autism biology.

To investigate this hypothesis, we first utilized a systems biological approach to determine the representation of high-confidence autism proteins and their interactors among empirically-derived cilia-related proteomes, identifying significant enrichment, particularly at more proximal regions of the ciliary basal body. We also observed that expression patterns of high-confidence autism genes tend to be significantly correlated with canonical cilia marker genes. Next, we conducted a localization screen of the 30 autism proteins with the strongest statistical association with autism and identified 12 that localize to both primary cilia on neurons as well as on motile cilia on epithelial cells. These proteins include both genes categorized as gene expression regulators as well as those characterized as canonical neuronal communication proteins such as SYNGAP1 and PSD95, suggesting that these proteins are pleiotropic and share a role at the cilium. Indeed, we observed that chromatin regulators CHD8, CHD2, and POGZ, as well as synaptic protein SYNGAP1, not only localized to cilia, but were required for their formation. Further, autism-associated patient missense variants in *SYNGAP1* mislocalized the protein away from cilia, and humans with pathogenic *de novo* variants in *SYNGAP1* produced significantly lower nasal nitric oxide gas, a canonical clinical biomarker for cilia function.

In sum, this work identifies convergence of autism proteins at cilia, spanning functional annotations, reframing their potentially relevant biology to autism beyond gene expression regulation and neuronal communication. This reconceptualizes our understanding of “chromatinopathies” and “synaptopathies,” highlighting their common biology with ciliopathies and the limitations of relying on gene ontologies in the pursuit of identifying relevant molecular mechanisms underlying a particular trait or condition. These findings provide new mechanistic insights into autism, establish the relevance of ciliary biology, and generate additional avenues for biomarker and therapeutics development.

## Results

### Autism proteins are enriched within ciliary proteomes

To begin to quantify the extent of convergence between ciliary biology and autism biology, we first sought to identify whether protein-protein interaction networks built from ciliary proteins (O’Neill et al. 2022; Gheiratmand et al. 2019) are enriched for high-confidence autism proteins. We previously demonstrated (Lasser et al. 2023a) that proteomes built from proteins of centriolar satellites and centrosomes (O’Neill et al. 2022; Gheiratmand et al. 2019), two structures required for organizing, building, and maintaining the cilium (Odabasi et al. 2019; Hall et al. 2023; Zhang and Mitchell 2015; Derderian et al. 2023), are enriched for high-confidence autism proteins FDR<0.1 (Fu et al. 2022) (summarized in **Fig. 1A**). Recently, a protein-protein interaction network derived from 100 high-confidence autism protein baits identified over 1,800 novel high-confidence protein-protein interactions (Autism PPI Network) (Wang et al. 2024). Therefore, based on the hypothesis that the interactors of autism proteins will also intersect ciliary biology, we tested whether the ciliary proteomes are enriched for proteins within the Autism PPI Network and observed significant overrepresentation (**Fig. 1A**, hypergeometric test, p < 0.05). This enrichment persists even when the high-confidence autism proteins used as baits are removed from the analysis (**Fig. 1A, Table S1**), suggesting that the overlap between autism and ciliary biology extends beyond genes directly identified in large-scale sequencing efforts. As these are microtubule-associated proteomes, we investigated whether our results might be non-specific such that any cytoskeletal proteome would show a similar enrichment. However, a proteome derived from actin-based structures (Thul et al. 2017) was not enriched for autism proteins or the Autism PPI Network (**Fig. 1A, Table S1**). To further evaluate the statistical significance of these findings, we conducted permutations tests with 200,000 iterations for each pairwise comparison of proteomes and observed significant enrichment of autism proteins in ciliary proteomes for all but 1 comparison, but never within the actin proteome (**Fig. S1, Table S1**).

**Figure 1:**
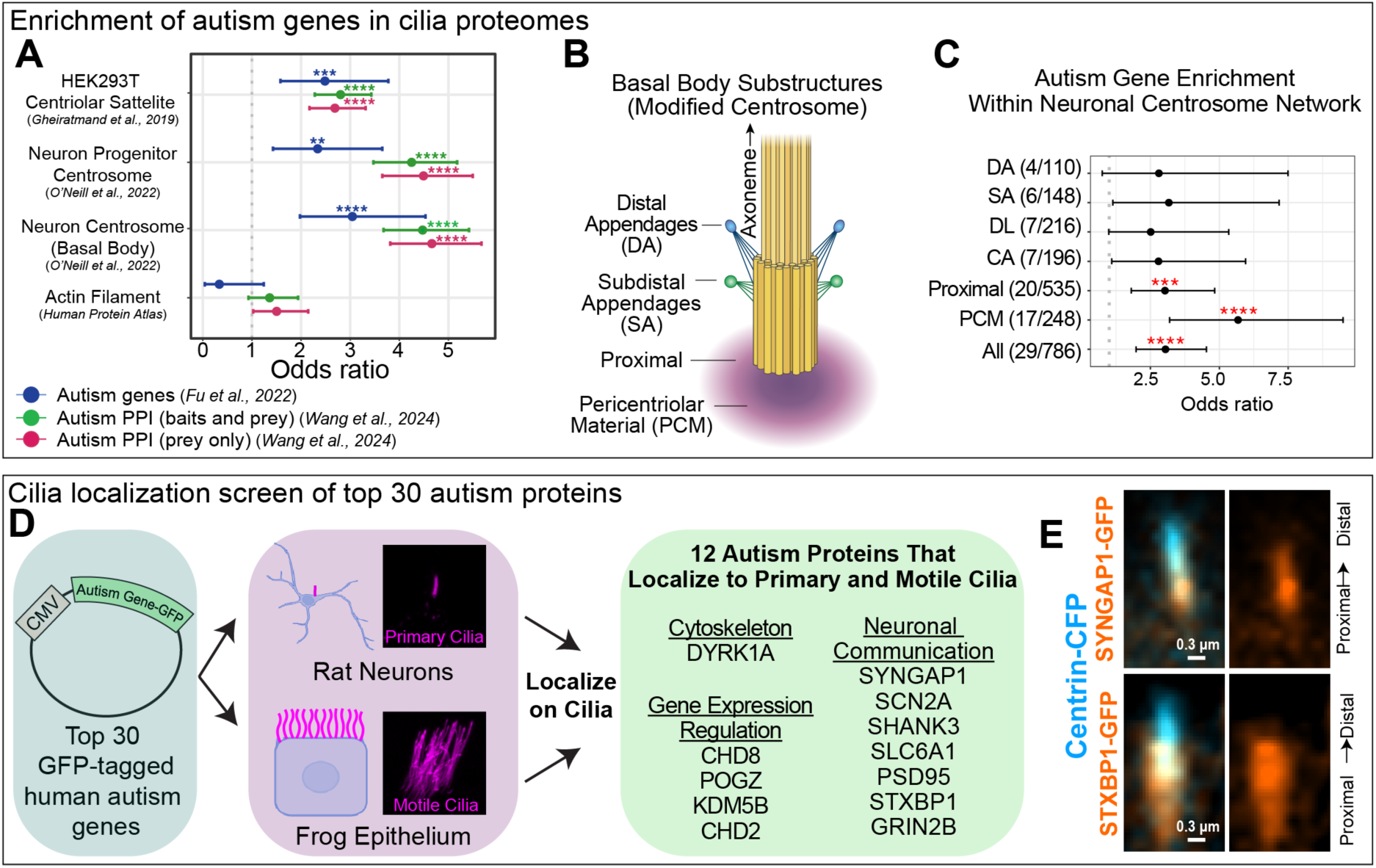
Autism proteins are enriched at cilia. (A) Enrichment of autism genes (Fu et al. 2022) and/or their protein-protein interactors (Wang et al. 2024) in three sets of cilia-related protein-protein interaction networks and one actin proteome (O’Neill et al. 2022; Gheiratmand et al. 2019; Thul et al. 2017). (B) Schematic of a ciliary basal body. (C) Enrichment of autism genes (Fu et al. 2022) within sub-ciliary structures of the post-mitotic neuronal centrosome protein-protein interaction network (O’Neill et al. 2022). DA: Distal Appendages (ODF2); SA: Subdistal Appendages (CEP170, ODF2); DL: Distal Lumen (POC5); CA: Cartwheel Assembly (CEP135); Proximal: proximal centriolar proteins (CEP62, CEP152); PCM: Pericentriolar Material (CEP192, CEP152, CDK5RAP2) from (O’Neill et al. 2022). Pericentriolar material (PCM) core protein (CENTROB) had 0/24 overlap and distal tip protein (CP110) had 0/85 overlap, so they were not included in the graph. (D) Schematic depicting localization screen in which 30 autism proteins with the strongest association with autism (FDR < 10^-6^, (Fu et al. 2022)) were tagged with GFP and overexpressed in motile cilia of Xenopus epidermis in vivo or primary cilia rat striatal neurons in vitro. Proteins listed were found to localize to cilia assessed by observation of GFP-tagged protein or endogenous antibody stain or both. (E) Human GFP-tagged SYNGAP1, STXBP1 localize proximally at the basal body (labeled by centrin-CFP, cyan), imaged live. See also Fig. S1-3 and Tables S1-3.

Next we sought to determine whether any particular sub-ciliary structures were predominantly responsible for the observed enrichment. We focused on the neuronal centrosomal proteome, given the strong evidence for significant co-expression of autism genes in neurons (Willsey et al. 2013; Parikshak et al. 2013), and since centrosomal structures in post-mitotic cells will be more directly related to ciliary biology as these cells lack mitotic spindles. We observed that the enrichment was driven by interactions with proteins known to localize to the proximal centrosomal and pericentriolar material (PCM) regions of the basal body, the more proximal regions of cilia, rather than by interactions with more distal proteins (**Fig. 1B-C, Table S1**).

We then tested whether gene expression of high-confidence autism genes correlated with expression of the canonical markers of primary cilia (*ARL13B*) and motile cilia (*FOXJ1*) in human single cell RNA sequencing datasets. We observed that 162 of the 255 genes identified as high-confidence autism genes (FDR < 0.1) in a recent large-scale genomics study of autism (Fu et al. 2022) show significant correlation with *ARL13B* and/or *FOXJ1* (Pearson correlation, corrected p < 0.05) in epithelial cells from the human Tabula Sapiens dataset (Tabula Sapiens Consortium* et al. 2022), including the classic cilia determinant *RFX3* (Bonnafe et al. 2004), (**Fig. S1, Table S2**). One of the genes most highly correlated with *FOXJ1* expression is the neuronal communication gene *SYNGAP1* (**Table S2**). Additionally, 101 of these 255 high-confidence autism genes are significantly correlated with *ARL13B* (Pearson correlation, corrected p < 0.05; *FOXJ1* was not analyzed here since few cells express it) in a human brain-specific dataset (Siletti et al. 2023) (**Fig. S1, Table S2**). These results suggest that high-confidence autism proteins are related to ciliary biology. Notably, the autism genes and proteins that are enriched in ciliary proteomes and correlated with canonical cilia markers include many proteins categorized as gene expression regulators or neuronal communication related proteins, suggesting that these broad functional categories do not fully capture the diversity of biological roles of these genes.

### Autism proteins of diverse functional annotations localize to cilia

Given a striking overlap between autism-associated proteins and protein interaction networks derived from proteins with known ciliary localizations, we next tested the hypothesis that autism-associated proteins commonly localize to cilia. Cilia are typically classified as either primary (like those in neurons) or motile (like those in the respiratory tract). While many ciliary proteins localize and function at both types of cilia, there are many exceptions (Hansen et al. 2024; Doerner et al. 2015), therefore we elected to test localization of autism proteins at both types of cilia. To do this, we selected the 30 proteins with the strongest statistical evidence of association with autism (FDR < 10^-6^, (Fu et al. 2022)) and expressed GFP-tagged human versions of these proteins in an *in vitro* model of neuronal primary cilia, primary rat embryonic striatal neurons in culture (Fagan et al. 2024) and in an *in vivo* model of motile cilia, the *Xenopus* embryonic epidermis (Werner and Mitchell 2013; Walentek 2021; Walentek and Quigley 2017; Rao and Kulkarni 2021) (**Fig. 1D**). We observed that 15 of these 30 autism proteins localize to neuronal primary cilia, 18 to motile cilia, and 12 to both primary neuronal cilia and motile cilia (**Fig. 1D, Table S3).** As expected in light of previous work from our group and others (Willsey et al. 2020; Lee et al. 2022), one of these proteins that localized to cilia was the cytoskeletal protein DYRK1A. The other 11 proteins that localized to both types of cilia have been functionally categorized as regulators of gene expression or as involved in neuronal communication (**Fig. 1D**). As in our prior functional work, this suggests that these proteins may have cellular functions beyond existing functional categorizations, and both categories will be explored in greater detail below (**Fig. 2-3**). Echoing our *in silico* work above, we imaged 3 proteins (SYNGAP1, STXBP1, and KMT5B) live at high resolution and observed proximal basal body localization relative to the basal body marker Centrin for all 3 (**Fig. 1E, S2**). Furthermore, of these 12 proteins that localize to cilia, detailed patient medical record review data from Citizen Health (Citizen) were available for 5 of these genes (*CHD2*, 42 patients, *SYNGAP1*, 188 patients; *SLC6A1*, 49 patients; *SCN2A*, 62 patients; *STXBP1*, 81 patients). Our analysis of these data revealed evidence of co-occurring cilia-related conditions among all cohorts (**Fig. S4, S5, Table S4**), including structural heart defects, craniofacial defects, and digit patterning defects, supporting the hypothesis that ciliary biology is disrupted in these patients and that variants in these genes can cause the cilia-related co-occurring conditions present in autism more generally. Together, these results support the hypothesis that high-confidence autism proteins spanning functional categorizations converge at cilia, based on both mass-spectrometry-based proteomic network analysis and microscopy-based localization screening.

**Figure 2:**
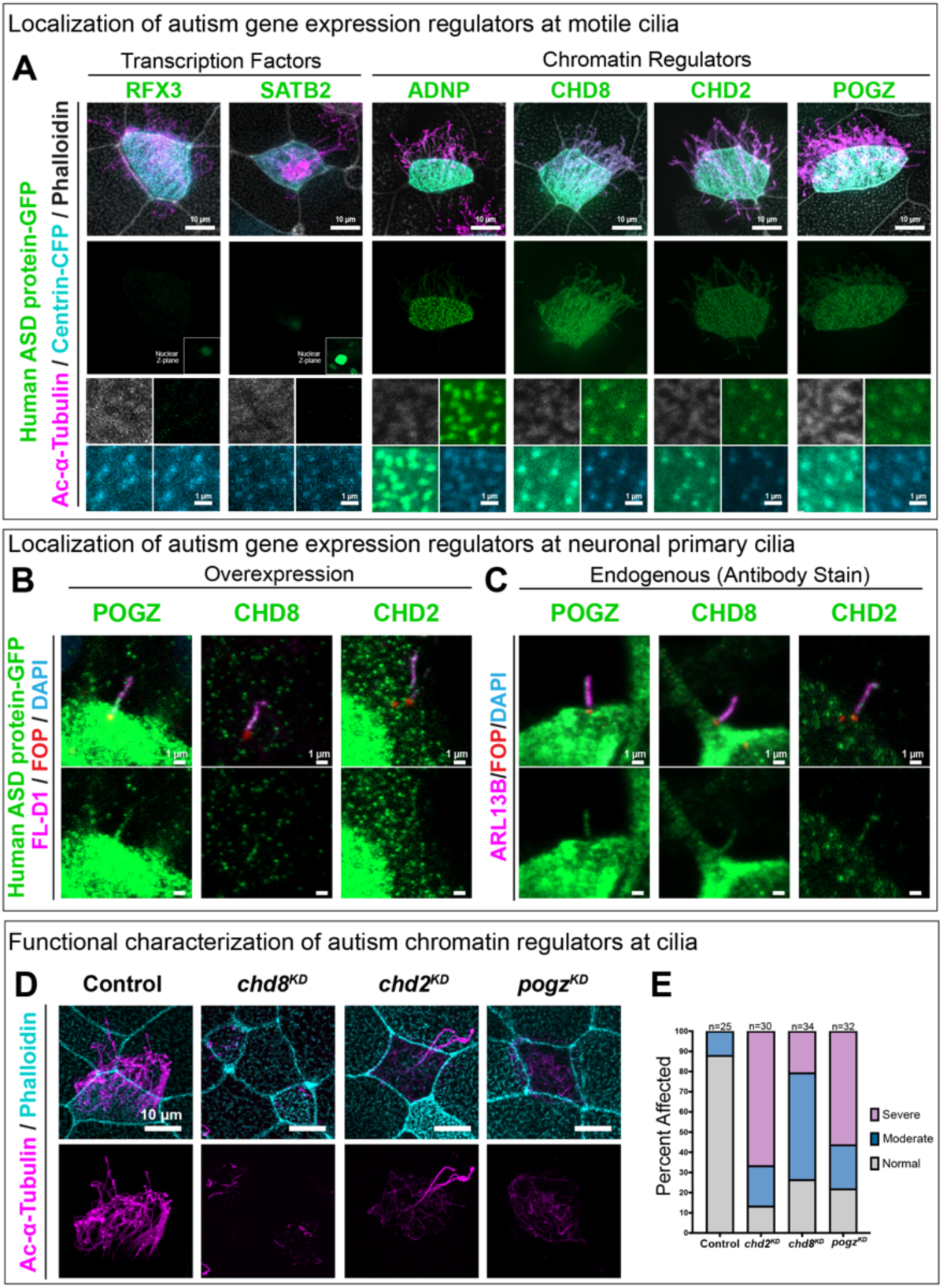
Autism-associated chromatin regulators localize to and function at cilia. (A) Human GFP-tagged constructs for autism-associated transcription factors SATB2 and RFX3 (green) localize to the nucleus (inset, deeper z-plane) and do not localize to motile cilia, while GFP-tagged constructs for autism-associated chromatin regulators ADNP, CHD8, CHD2, and POGZ do localize to motile cilia (labeled by acetylated alpha-Tubulin, magenta; basal bodies labeled by Centrin-CFP, cyan) when expressed in the X. laevis embryonic epidermis. Bottom panel shows basal bodies, actin, and autism-associated proteins in greater detail (actin network labeled by phalloidin, gray). (B) Human GFP-tagged constructs for POGZ, CHD8, and CHD2 (green) localize to primary cilia (cilia marked by Flag-tagged D1 receptor, magenta; basal bodies labeled by FGFR1OP antibody, red) when expressed in primary rat striatal neurons. (C) Endogenous staining for POGZ, CHD8, and CHD2 (green) in primary cilia (labeled by ARL13B, magenta; basal bodies labeled by FGFR1OP, red) in primary rat striatal neurons. (D) Morpholino-mediated knockdown (KD) of chd8, chd2, or pogz in X. tropicalis results in defects in cilia (labeled by acetylated alpha tubulin, magenta) on multiciliated cells of the embryonic epidermis. (E) Quantification of the data shown in B, with cilia.

**Figure 3:**
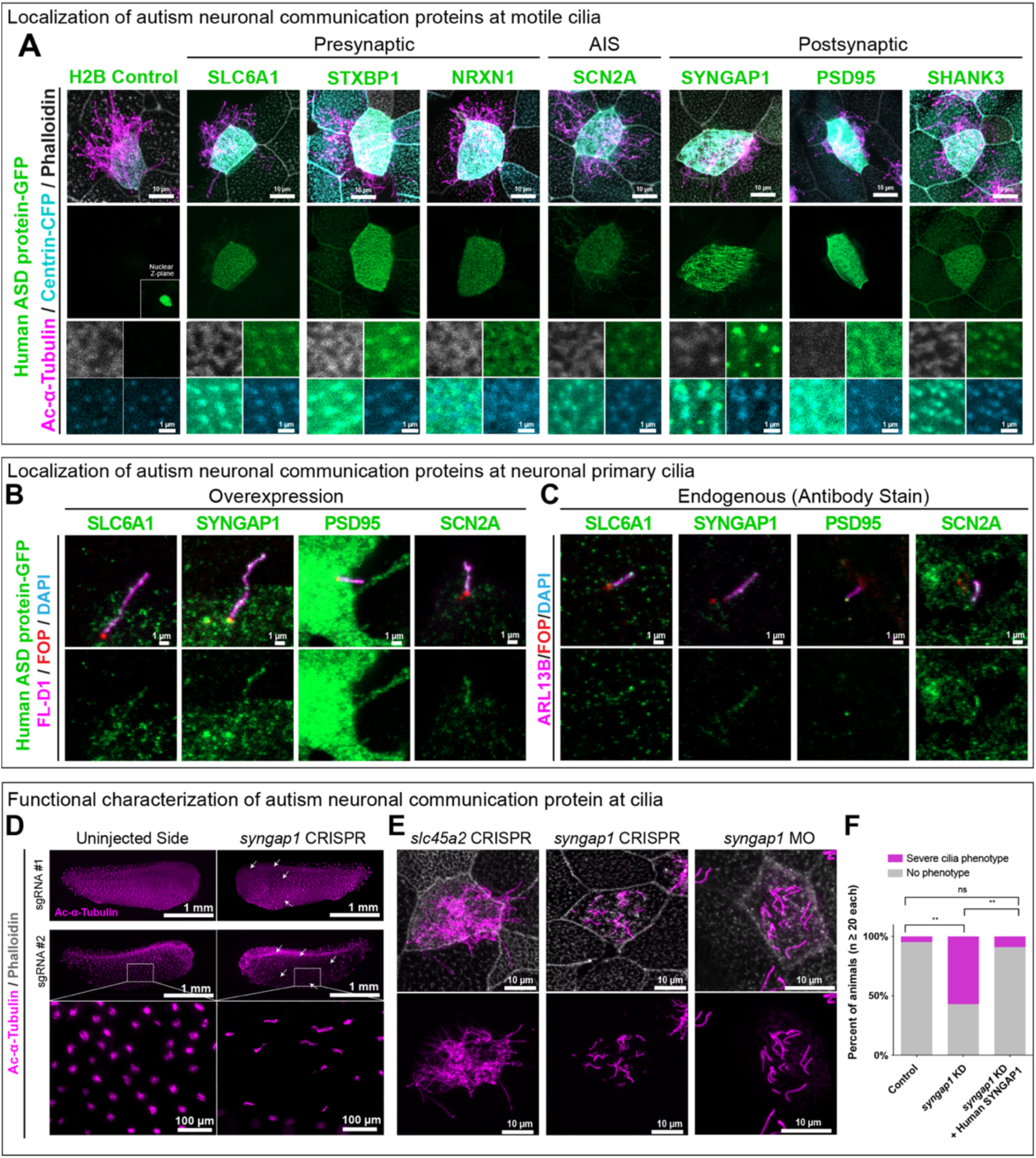
Autism-associated neuronal communication proteins localize to and function at cilia. (A) Human GFP-tagged constructs for autism-associated ‘neuronal communication’ proteins (green) known to localize across neuronal subcompartments— presynaptic density, axon initial segment (AIS), and postsynaptic density—localize to cilia (axonemes labeled by acetylated alpha-Tubulin, magenta; basal bodies labeled by centrin-CFP, cyan) when expressed in X. laevis. NRXN1 localizes to the actin network surrounding cilia (labeled by phalloidin, gray). Bottom panel shows basal bodies and autism proteins in detail. H2B-GFP control localizes to the nucleus, in a deeper Z-plane as indicated in the inset “Nuclear Z-plane.” (B) Human GFP-tagged constructs for SLC6A1, SYNGAP1, PSD95, and SCN2A (green) localize to primary cilia (cilia labeled by Flag-tagged Dopamine D1 receptor, magenta; basal bodies labeled by FGFR1OP, red) when expressed in primary rat striatal neurons. (C) Endogenous staining for SLC6A1, SYNGAP1, PSD95, and SCN2A in primary cilia (labeled by ARL13B, magenta; basal bodies labeled by FGFR1OP, red) in primary rat striatal neurons. (D,E) Loss of syngap1, via CRISPR (syngap1^CR^) or morpholino-mediated knockdown (syngap1^KD^), results in cilia defects (labeled by acetylated alpha-Tubulin, magenta) and disrupts the apical actin network (labeled by phalloidin, gray) in the X. tropicalis multiciliated epidermis. (F) Quantification of syngap1^KD^ displayed in E as a percentage of embryos with no cilia phenotype vs. with a moderate or severe cilia phenotype. Fisher’s exact test (two-sided) followed by Pairwise Fisher’s Exact tests were used to calculate significance using raw counts of individual embryos. ** = p < 0.01, ns = no statistically significant difference. See also Fig. S8-11 and Tables S3 and S5.

### Autism gene expression regulators localize to and function at cilia

Autism-associated proteins classified as functioning in gene expression regulation are largely transcription factors or chromatin regulators (Sestan and State 2018; Satterstrom et al. 2020; Willsey et al. 2022). As expected, transcription factors such as RFX3 and SATB2 localized to the nucleus, but not to the cilium (**Fig. 2A**). Indeed, among the autism proteins tested here that do not localize to cilia, genes that are functionally annotated as transcription factors make up the largest proportion (**Fig. S3**). In contrast, we observed eight autism proteins functionally annotated as chromatin regulators localizing to motile cilia and four of these eight also localizing to neuronal primary cilia, as assessed by overexpression of GFP-tagged protein (**Fig. 2A-C, S6, Table S3).** This is consistent both with our recent work demonstrating that five high-confidence autism-associated chromatin regulators not only localize to the nucleus, but also to microtubules of the mitotic spindle (Lasser et al. 2023), as well as the known roles of other chromatin regulators at the cilium including HDAC6, WDR5, and CHD4 (Hubbert et al. 2002; Ran et al. 2015; Kulkarni et al. 2018; Marley and von Zastrow 2012; Teerikorpi et al. 2024). Given that overexpression of GFP-tagged proteins can lead to improper protein localization, we confirmed these localizations for the 3 chromatin regulators that have not otherwise been shown to function at cilia (POGZ, CHD8, CHD2) using antibodies that we orthogonally validated (**Fig. 2C, S7**). Since cilia, like mitotic spindles, are microtubule-based structures, these findings suggest a potentially broader role for autism chromatin regulators in microtubule biology. Consistent with this notion, the autism-associated chromatin regulator *KMT5B* has recently been shown to be required for ciliogenesis in *Xenopus* (Angerilli et al. 2023), and ADNP is also known to stabilize microtubules directly (Gozes et al. 2018).

While it is striking that many autism chromatin regulators localize to ciliary structures, it is possible that they do not function at cilia endogenously. To investigate this, we selected 3 of these proteins (CHD2, CHD8, and POGZ) that have no known prior function at the cilium and depleted them individually during embryogenesis in the diploid species *X. tropicalis*, and, like KMT5B (Angerilli et al. 2023), observed ciliogenesis defects in the multiciliated epidermis (**Fig. 2D, S7**). Together, these results demonstrate that multiple high-confidence autism proteins annotated as chromatin regulators can localize to cilia and in some cases are required for their formation.

### Autism neuronal communication proteins localize to cilia

We recently described, using a *Xenopus* model, that several high-confidence autism genes with canonical roles in neuronal communication, including the gene *SYNGAP1*, are required for forebrain neurogenesis prior to synaptogenesis (Willsey et al. 2021). This finding has since been replicated in human cortical organoids and rodents (Birtele et al. 2023). We also previously showed that the autism associated sodium channel, SCN2A, also classified into neuronal communication, is part of a protein interaction network at the interface of autism and congenital heart disease, with *in vivo* perturbation causing neurogenesis and structural heart defects (Rosenthal et al. 2021; Willsey et al. 2021), common outcomes of ciliary dysfunction. Together, these results suggested that *SYNGAP1*, *SCN2A*, and potentially other autism genes included in the neuronal communication group have functional roles beyond neurotransmission, including during neurogenesis and heart development, processes that depend on cilia (Park et al. 2019)

Among the 30 proteins with the strongest statistical evidence for association with autism, nine are annotated as having a role in neuronal communication (Satterstrom et al. 2020). We observed seven of these proteins localizing to motile and primary cilia, either by expression of the GFP-tagged human protein and/or by endogenous antibody staining (**Fig. 1D, 3A-C, S8**). These proteins have been otherwise characterized to function across three neuronal subcellular compartments: the postsynaptic density, the axon initial segment, and the presynaptic density. GFP-tagged STXBP1 and GRIN2B strongly localized to ciliary basal bodies of motile cilia (**Fig. 3A, S8**), but we were unable to achieve expression of these transgenes in neurons. However, by endogenous antibody staining, both STXBP1 and GRIN2B localized to primary cilia in cultured neurons (**Fig. S8, Table S3**), so we classified them as localizing to both kinds of cilia. One presynaptic neuronal communication protein, NRXN1, did not localize to motile cilia, but rather to the actin matrix that stabilizes motile cilia (**Fig. 3A**).

We also confirmed ciliary localizations for SYNGAP1, STXBP1, PSD95, SCN2A, SLC6A1, and SHANK3 by antibody staining in rat brain cells (**Fig. 3C, S8C, S9A**), using antibodies that have been empirically tested for specificity and/or previously published (Bayés et al. 2012; Sumathipala et al. 2023; Birtele et al. 2023; Mastro et al. 2020; Unterauer et al. 2024; Puntman et al. 2021; Cijsouw et al. 2014; Wan et al. 2021; Okuzono et al. 2023; Ferhat et al. 2023; Landry et al. 2023; Verret et al. 2012). Together, these results suggest that autism proteins with annotated functions in neuronal communication also localize to cilia and therefore may share a previously unrecognized common function at this structure.

### Neuronal communication protein SYNGAP1 functions in cilia formation

Many of these autism-associated ‘neuronal communication’ proteins that localize to cilia are ion transporters or receptors. Therefore, we anticipate that these proteins may function in ciliary signaling rather than in cilia formation. However, SYNGAP1 is a GTPase activating protein with known roles regulating Rac signaling and downstream actin dynamics (Carlisle et al. 2008; Su et al. 2019), which are critical for cilia formation (Hoffman and Prekeris 2022; Smith, Lake, and Johnson 2020; Kim et al. 2015). Consequently, we hypothesized that SYNGAP1 may have a role in cilia formation.

Because our localization screen required overexpression of the interrogated protein, which can lead to non-physiological localizations, we sought to confirm the endogenous localization of SYNGAP1 across multiple species and ciliated cell types, including neurons, epidermis, and respiratory tract. We observed endogenous expression and localization of SYNGAP1 in frogs, mice, rats, and human cells using RNA in situ hybridization, antibody staining, and/or available scRNA-Sequencing datasets (**Fig. 3, S9-S11**). In fact, *SYNGAP1* is expressed in human multiciliated cells (*FOXJ1*+) at similar levels as in neurons (**Fig. S11**). Next we aimed to test whether SYNGAP1 is required during cilia formation. To do this we perturbed *syngap1 in vivo* in *X. tropicalis* both by CRISPR-mediated mutagenesis and by morpholino-based translation-blocking. In both cases, we observed ciliogenesis defects in the embryonic epidermis (**Fig. 3D-F**). Supplying human SYNGAP1-GFP rescued the morpholino perturbation (**Fig. 3D-F, Table S5**). As expected given its published role in actin regulation (Carlisle et al. 2008; Su et al. 2019), we observed defects in cortical actin organization (**Fig. 3E**). Together, this work demonstrates that SYNGAP1 is endogenously expressed in a wide variety of ciliated cell types across multiple species, localizes to diverse cilia, and is required, at least in some contexts, for cilia formation.

### Patient-derived SYNGAP1 variants mislocalize away from cilia

Next we sought to test whether autism-associated missense variants (Fu et al. 2022) affect ciliary localization of human SYNGAP1 protein. Two of the five autism-associated, likely pathogenic missense variants observed in Fu et al. (2022) (Fu et al. 2022) map within or near the protein’s annotated pleckstrin homology (PH) domain (**Fig. 4A**). PH domains are known to influence membrane-targeting (Singh et al. 2021; Vonkova et al. 2015; Lemmon et al. 1997) and, since cilia are membrane-bound structures, we hypothesized that these missense variants may influence ciliary localization. While the non-mutant human sequence localized to ciliary basal bodies and the base of axonemes when expressed in *Xenopus* (**Fig. 4B**), the two autism-associated missense variants (S140F and C233Y) showed altered ciliary localization, with an S140F mutation leading to a reduction at ciliary basal bodies and axonemes as well as to mislocalization of the mutated protein to the plasma membrane (**Fig. 4B-C**). A C233Y mutation similarly led to a reduction of protein at basal bodies and axonemes, but with an additional accumulation in liquid-like aggregates within the cytoplasm, and no plasma membrane mislocalization (**Fig. 4B, D**). These results demonstrate that patient-derived missense variants have the potential to cause SYNGAP1 to mislocalize away from the cilium.

**Figure 4:**
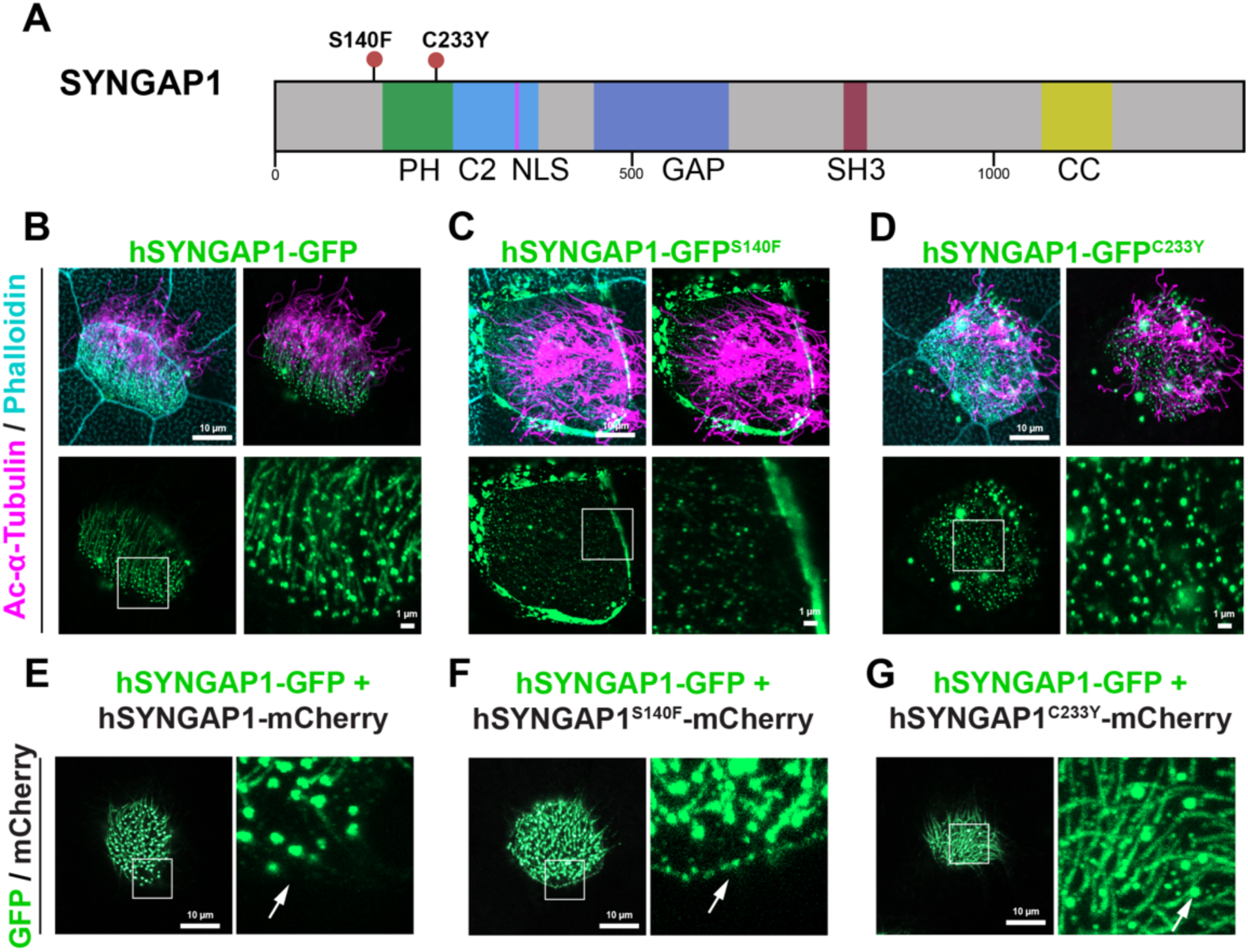
Patient-derived SYNGAP1 missense variants mislocalize away from cilia. (A) Schematic of human SYNGAP1 protein (isoform alpha-2) with functional domains and locations of two autism-associated (Fu et al. 2022) patient-derived missense B variants annotated. (B-D) GFP-tagged human SYNGAP1 (green) localizes to the cilium (basal bodies labeled by centrin-CFP, cyan; axonemes labeled by acetylated alpha-Tubulin, magenta), while autism-associated variant SYNGAP1S140F often localizes away from the cilium at the membrane and SYNGAP1C233Y often localizes away from the cilium into aggregates. Actin network labeled by phalloidin (cyan). (E-G) GFP-tagged human non-mutant SYNGAP1 (green) localizes to the cilium when co-expressed with mCherry-tagged human non-mutant SYNGAP1 (E), but often localizes away from the cilium and to the membrane when co-expressed with mCherry-tagged autism-associated variant SYNGAP1S140F (F) or into aggregates when co-expressed with mCherry-tagged autism-associated variant SYNGAP1C233Y (G). mCherry proteins in white in the left panels. Right panels in 4G-I show GFP-tagged human non-mutant SYNGAP1 alone at higher magnification.

Of note, we used *SYNGAP1* isoform alpha-2 (**Fig. 4A**) for these experiments as this is the only C-terminal isoform that contains all reported autism-related likely pathogenic variants (Fu et al. 2022). This isoform, unlike alpha-1, lacks a C-terminal PDZ-binding motif but retains the coiled-coil-like domain that is considered important in the protein’s trimerization (Zeng et al. 2016). Since this isoform retains the coiled-coil-like domain, we hypothesized that these mutant proteins may still trimerize with non-mutant SYNGAP1 protein and cause it to mislocalize as well. Therefore, we tagged the missense variants of *SYNGAP1* with mCherry and assessed whether presence of the variant protein could alter the localization of a GFP-tagged non-mutant SYNGAP1 protein by co-expressing them. Indeed, for both variants, the GFP-tagged non-mutant protein mislocalized along with the missense variant protein (**Fig. 4E-G**). Together, these results show that select missense variants in SYNGAP1 can cause protein mislocalization away from the cilium in *Xenopus*.

### Reduced ciliary biomarker levels in affected *SYNGAP1* individuals

These findings so far suggest that *SYNGAP1*-related disorders may be considered ciliopathies. Consequently, we sought to directly test whether individuals with genetic variants in *SYNGAP1* show altered biomarkers of ciliary function. One canonical ciliary functional biomarker is nasal nitric oxide (nNO), which is used to diagnose the ciliopathy primary ciliary dyskinesia (PCD) (Leigh et al. 2013; Shapiro et al. 2018; Beydon et al. 2023). NO gas is produced by nasal multiciliated cells (which express both *FOXJ1* and *SYNGAP1*) (**Fig. 5A, S1, S11, Table S2**), and patients with cilia defects have characteristically low nNO values (Narang et al. 2002; Wodehouse et al. 2003) (**Fig. 5B**). We therefore measured nNO of 24 individuals with *de novo* variants in *SYNGAP1* as well as a matched unaffected immediate family member. Individuals with *SYNGAP1* variants had significantly lower nNO values compared to their family members (p = 0.0015, one-sided Wilcoxon matched-pairs signed rank test, **Fig. 5C**), which was not due to differences in sex or age (**Fig. 5D, S12A-B**). Further, the effect was statistically significant when compared unpaired (p = 0.0024, one-sided Mann-Whitney rank sum test, **Fig. S12F**). Splitting out variant classes, data from individuals with PTVs alone (n = 21 patients) are significant (p = 0.0051, one-sided Wilcoxon matched-pairs signed rank test, **Fig. S12C**), and values from individuals with missense variants (n = 2) or a VUS (n = 1) have consistent directions of effect (**Fig. S12D-E**).

**Figure 5:**
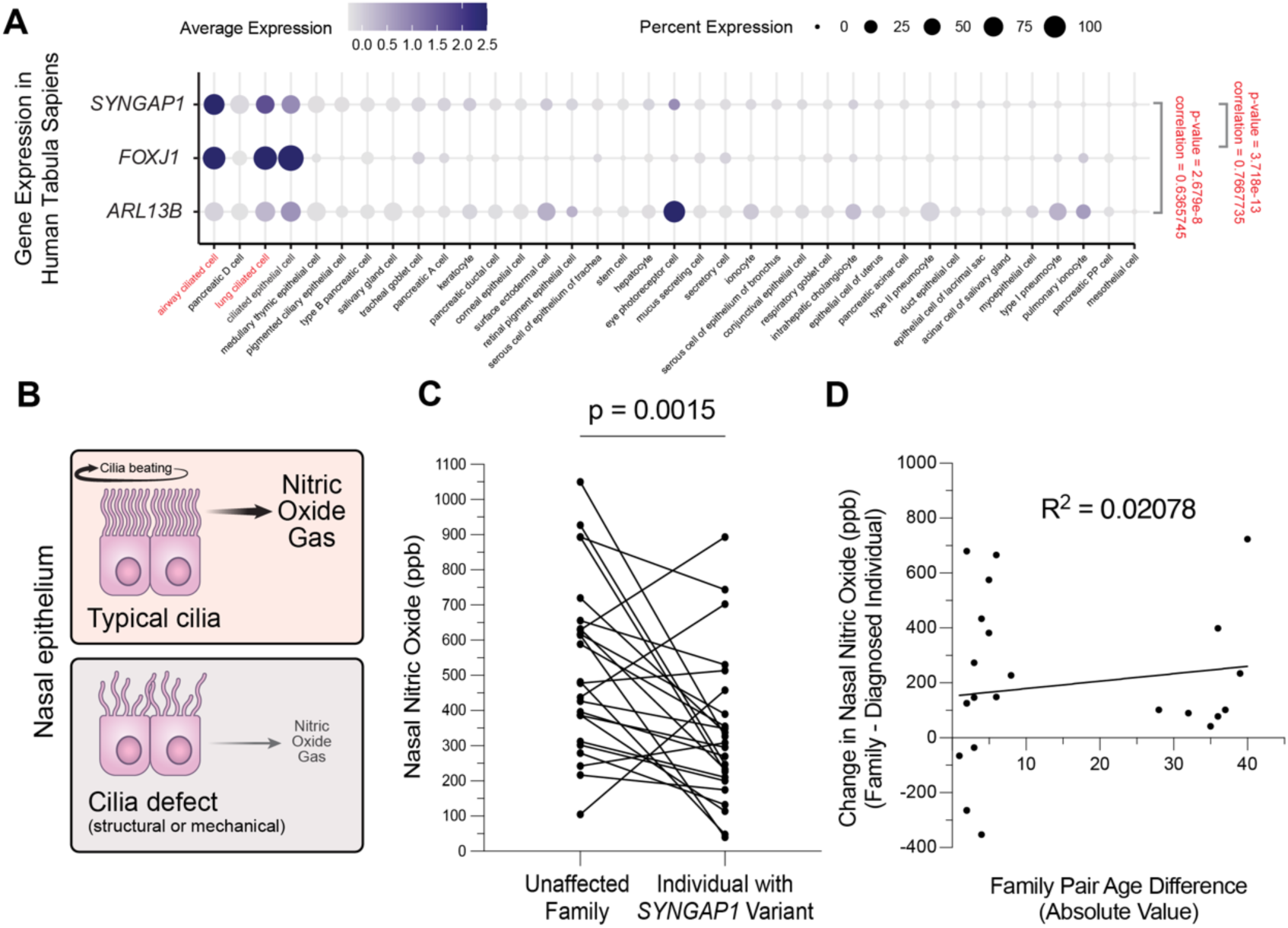
Ciliary functional biomarker nasal nitric oxide is reduced in SYNGAP1 patients. (A) SYNGAP1 is highly expressed in human airway ciliated cells and is significantly correlated with cilia markers FOXJ1 and ARL13B in the Tabula Sapiens epithelial dataset. (B) Nasal airway ciliated cells produce nitric oxide gas, but in the presence of cilia defects, this is greatly reduced. (C) Patients with de novo variants in SYNGAP1 have significantly lower nasal nitric oxide levels (p = 0.0015, one-sided Wilcoxon matched-pairs signed rank test, NIOX VERO nNO machine, tidal breathing method) compared to matched immediate family members (sibling or parent). (D) There is no relationship between family pair age difference and change in nitric oxide value (Simple Linear Regression, R^2^ = 0.02078). ppb = parts per billion. See also Fig. S12.

Two patients (p.Gln756* and p.His1203Thrfs*3) had nNO values lower than the current established diagnostic nNO cutoff for PCD (< 30 nL/min for tidal breathing method, which is < 100 ppb on this machine) (Beydon et al. 2023; Paternò et al. 2023), and no unaffected family members were below this value. It is not surprising that only a few patients met this cutoff, as these *SYNGAP1* variants are *de novo* heterozygous variants, while PCD is most commonly due to homozygous loss of function variants (Horani and Ferkol 2021). Thus, these results are consistent with ciliary dysfunction being a component of *SYNGAP1*-related disorders and motivates future work on similar patient cohorts for high confidence autism genes expressed in nasal multiciliated cells with high prevalence of cilia-related co-occurring conditions.

## Discussion

Despite the knowledge that autism commonly co-occurs with many disorders with known ciliary contributions, the role of autism genes in ciliary biology and the possibility that ciliary biology underlies a subset of autism has not been systematically explored. Here we show extensive convergence of high-confidence autism proteins at the cilium, a structure required for neurogenesis, dorsal/ventral forebrain patterning, and neuronal activity (Hasenpusch-Theil and Theil 2021; Tereshko et al. 2021; Park et al. 2019)—all processes that have repeatedly been implicated in autism biology (Willsey et al. 2022; Sun et al. 2024; Packer 2016).

While prior work by our group has demonstrated that ciliary biology is relevant to several high-confidence autism genes (Lasser et al. 2023; Willsey et al. 2020, 2018; Sun et al. 2024; Teerikorpi et al. 2024; McCluskey et al. 2024), this work more directly establishes a link between ciliary biology and large-effect, high-confidence autism genes, building on current knowledge that several “syndromic” autism proteins such as TSC1/2, FMRP, and MECP2 also localize to cilia and are required for cilia formation, maintenance, and/or signaling (Di Nardo et al. 2020; Karalis et al. 2022; Lee et al. 2020; Frasca et al. 2020; Hao et al. 2021). Given the great degree of coincidence between cilia-related disorders and autism, including congenital heart disease, hydrocephalus, asthma, and blindness (Sigmon et al. 2019; Rosenthal et al. 2021; Al-Beltagi 2021; A. J. Willsey et al. 2018; Duy et al. 2022; Jin et al. 2017; Pili et al. 2021), these findings suggest that the role of cilia should be considered further in autism and in commonly co-occurring and oft-disabling medical conditions like epilepsy, gastrointestinal disorders, congenital heart disease, respiratory issues, sleep disorders, and sensory sensitivities (Al-Beltagi 2021; Danesh et al. 2021). Indeed, a survey of autistic individuals and their family members revealed that understanding commonly co-occurring conditions is of highest importance to autistic people (Frazier et al. 2018).

This work highlights the well-known limitations of relying on functional annotations in the pursuit of identifying relevant molecular mechanisms underlying a particular condition or trait. Further, it demonstrates that even when one function of a gene has been well-established, it is still possible that the gene has additional relevant functions. In the case of autism genetics, the emphasis on “gene expression regulation” and “neuronal communication” may have prematurely limited the exploration of other plausible convergent molecular mechanisms. Indeed, here we demonstrate that autism-associated chromatin regulators also localize to microtubule-based ciliary structures, consistent with the work from our group and others that chromatin regulators influence both the tubulin and histone codes (Lasser et al. 2023; Park et al. 2016; Yokoyama et al. 2013; Sillibourne et al. 2007; Hubbert et al. 2002). The mechanism by which these proteins regulate ciliogenesis, whether in the nucleus, at the cilium, or both, will be an important future area of investigation. In addition, we provide evidence that genes with well-established roles at the synapse, like *SYNGAP1*, also localize to and function at the cilium. Consistently, SYNGAP1 was previously shown to interact with dopamine receptor 1 (Su et al. 2019), a known neuronal ciliary receptor (Domire et al. 2011; Stubbs et al. 2022; Leaf and Von Zastrow 2015; Marley and von Zastrow 2010). Similarly, ANK2, annotated in neuronal communication, was previously localized to cilia and shown to be required for ciliogenesis in human epithelial cells (Gupta et al. 2015). While this manuscript was in preparation, SLC6A1 and STXBP1 were reported in a preprint to localize to mouse neuronal primary cilia by proximity-ligation mass spectrometry (Loukil et al. 2023). Together, these results suggest that multiple autism proteins localize to cilia and therefore may share an additional function at this structure. It is also possible that this work underestimates the functional convergence of genes on the cilium, as genes may not localize to the cilium, but may still affect cilia function or cilia formation. For example, here we show that RFX3, a transcription factor, does not localize to cilia; however, it is well-established that RFX3 is a critical regulator of ciliogenesis (Choksi et al. 2014; Chen et al. 2018). Similarly, NRXN1 did not localize to cilia, but instead localized to the apical actin network surrounding basal bodies that stabilizes motile cilia and are essential for ciliogenesis (Sedzinski et al. 2017; Hoffman and Prekeris 2022; Sedzinski et al. 2016). Conversely, it is possible that some of these proteins that localize to cilia do not function there.

Many of these autism genes with roles in synaptic biology and in neurotransmission are also associated with intractable epilepsy, and genetic perturbations are often termed “synaptopathies” (Spoto et al. 2022). However, as cilia defects can cause intractable epilepsy (Reiter and Leroux 2017), this work suggests that ciliary dysfunction may contribute to the co-occurring seizures and provides an additional avenue for therapeutics development, particularly for these treatment-refractory epilepsies. Similarly, an exciting future opportunity lies in understanding the similarities between the cilium and the synapse. These subcellular structures are both membrane-bound projections that are rich in receptors, have diffusion barriers, active microtubule-based transport, signal through calcium, regulate neuronal excitability, and modulate downstream signaling pathways (Nechipurenko et al. 2013; Delling et al. 2013; Tereshko et al. 2021; Huang and Rasband 2018; Park and Leroux 2022). Understanding these structures’ similarities, differences, and intersections will be important future work, particularly as neuronal ciliary biology is explored deeper and direct interactions between neuronal cilia and synapses are elaborated (Sheu et al. 2022).

In conclusion, we have established through multiple lines of investigation in a range of model systems, as well as in humans, that many autism-associated proteins of diverse functional annotations localize to and function at the cilium, leading to a new and important area of investigation in the search for an actionable understanding of autism and related developmental disorders. Points of convergent biology hold promise for developing new biomarkers and for rational design of therapeutics that may be relevant to a wide range of individuals, potentially avoiding the concern that treatments will need to be developed for each individual large-effect gene. Therefore, a deeper investigation of the relative contributions of ciliary biology, gene expression regulation, and neuronal communication to the etiology of autism is a critical direction for future investigation.

## Acknowledgements

First and foremost, we thank the SYNGAP1 patients and their families for participating in research as well as the SynGAP Research Fund. We thank Citizen Health and all patients and families that participated. We thank: Aaron Marley and Mark von Zastrow for early preliminary SYNGAP1 and ANK2 localizations in IMCD3 cells; Nolan Wong and UCSF LARC for animal care; Milagritos Alva for lab maintenance; Ashley Clement, Gigi Lopez, Sonia Lopez, and Linda Chow for administrative support; Martin Kampann for plasmid phr_tet3g; Tom Nowakoski for generous use of his confocal microscope; Jeremy Reiter, Kunal Shroff, Yue Liu, Mia Konjikusic, Elena Turkalij, Angeline Chemel, Dylan Yep, Thi Nguyen, Gabriella Canales, Camille Derderian, and Thao Phan for advice and feedback; Sarah Pyle for graphic design work; the Berkeley Molecular Imaging Center (NIH grant S10OD025063) and Cherise Stanley for microscopy assistance; and Kari Herrington and Micaela Lasser at the UCSF Center for Advanced Light Microscopy for microscopy assistance. This work would not be possible without daily reference to the *Xenopus* community resource Xenbase (RRID:SCR_003280) and expertise and frog resources from the National Xenopus Resource (RRID:SCR_013731). HRW is a Chan Zuckerberg Biohub - San Francisco investigator.

## Funding

Overlook International Foundation (MWS)

National Institutes of Health Convergent Neuroscience Initiative Grant U01MH115747 (MWS, AJW)

National Institutes of Health Grant U01MH116487 (MWS, AJW) Chan Zuckerberg Biohub - San Francisco (HRW)

Coalition to Cure CHD2 (HRW)

The SynGAP Research Fund (HRW)

The Sorensen Foundation Fellowship in Child & Adolescent Psychiatry (BW)

National Institutes of Health DSPAN Award F99 NS139536 (EK)

## Author Contributions

Conceptualization: HRW, AJW

Formal Analysis: EK, MCL, BW, JS, AS, KEM, AE, HRW

Funding Acquisition: MWS, AJW, HRW

Investigation: EK, HRW, MCL, BW, JS, AS, KEM, EBader, AE, CB, OC, JA, JD, HJL, KAH, SW, EBrimble

Methodology: HRW, EK, AJW, BW, AE

Project Administration: HRW

Resources: JMG, AJW, MWS, HRW, DBK, AE

Supervision: HRW, AJW, MWS, BW, AE

Validation: EK, HRW, AE, CB, EBrimble

Visualization: EK, JS, BW, AS, KEM, HRW

Writing- original draft: HRW, EK

Writing- review & editing: EK, HRW, AJW, MWS

## Data and Materials Availability

PPI data are publicly available at (Gheiratmand et al. 2019; O’Neill et al. 2022; Wang et al. 2024) and autism genetic data are publicly available at (Fu et al. 2022). Citizen Health medical record data is available with IRB approval.

## Materials and Methods

### Autism protein enrichments among cytoskeletal proteomes

We evaluated three sets of autism-related genes:

1. “Autism genes”: 255 autism high-confidence risk genes identified in a recent WES/WGS omnibus study (FDR<0.1, (Fu et al. 2022)) are enriched for proteins associated with centriolar satellites (Gheiratmand et al. 2019) or centrosomes (O’Neill et al. 2022).
2. “Autism PPI (baits and prey)”: 1143 autism-associated proteins defined in an autism protein interaction network generated from by exogenously overexpressing 100 high-confidence autism risk genes (‘bait’) in HEK293T cells and identifying associated proteins (‘prey’) using affinity-purification mass spectrometry (Wang et al. 2024).
3. “Autism-PPI (prey only)”: 1043 autism-associated proteins that are not high-confidence autism risk genes (geneset 2 prey, excluding all bait).

We defined four sets of proteomes associated with various cytoskeletal components, including centriolar satellite, centrosome, or actin filaments and 8 sets of proteins associated with centrosomal subregions:

● HEK293T centriolar satellite (Gheiratmand et al. 2019): centriolar satellite proteome (HEK293T) generated using in vivo proximity labeling with 22 human satellite proteins in HEK293T cells. Interactors were downloaded from Gheiratmand et al. 2019 Table EV2 and gene names were updated using limma::alias2SymbolTable.
● NPC centrosome (O’Neill et al. 2022): centrosome-associated proteome generated by AP-MS of 10 centrosomal baits in human iPSC-derived NSCs (O’Neill et al. 2022). Interactors were downloaded from O’Neill et al. 2022 Table S7 (NSC_sig=1) and gene names were updated using limma::alias2SymbolTable.
● Neuron centrosome (and 8 subsets) (O’Neill et al. 2022): centrosome-associated proteome generated by AP-MS of 10 centrosomal baits in human iPSC-derived NSCs (O’Neill et al. 2022). Interactors were downloaded from O’Neill et al. 2022 Table S7 (Neuron_sig=1) and gene names were updated using limma::alias2SymbolTable. 8 subsets of interactors that associated with bait that localize to various centrosome subregions were defined (subregions based on O’Neill et al. 2022 Figure 1H, consisting of pericentriolar material, PCM core, proximal, cartwheel assembly, distal lumen, subdistal appendages, distal appendages, and distal tip). The 2 subregions without any overlap (PCM core 0/24, distal tip 0/85) were excluded from the graph since they did not have any overlap and had a relatively smaller number of interactors.
● Actin filament (Thul et al. 2017): Proteins that localize to actin filaments (https://www.proteinatlas.org/humanproteome/subcellular/actin+filaments) and the background set of 13,147 proteins whose subcellular localization was evaluated in the Human Protein Atlas (https://www.proteinatlas.org/search/has_protein_data_in%3ACell) were downloaded in May 2024. Gene names were updated using limma::alias2SymbolTable and genes without associated HGNC symbols were removed.

We evaluated whether the ASD-related genesets are significantly enriched in the centriole/centrosome-associated genesets. For the HEK293T centriolar satellite geneset (Gheiratmand et al. 2019), we defined the universe to be the intersection of genes evaluated in Fu et al. 2022 (Fu et al. 2022) and the HEK293T proteome (union of a set of proteins detected in HEK293T (Bekker-Jensen et al. 2017) and HEK293T centriolar satellite proteins (Gheiratmand et al. 2019). For the NPC and neuron centrosome genesets (O’Neill et al. 2022), we defined the universe to genes evaluated in Fu et al. 2022 (Fu et al. 2022) as the NPC and neuron proteomes were not reported in this study (O’Neill et al. 2022). For the actin filament geneset, we defined the universe to be the intersection of genes evaluated in Fu et al. 2022 and genes whose protein products were evaluated for subcellular localization in the Human Protein Atlas (Thul et al. 2017). We conducted Fisher’s exact tests (two-sided) using a 2×2 contingency table using the variables ASD-associated gene (yes/no) and proteome geneset (yes/no). We corrected p values for multiple hypothesis testing (p.adj = pval * # genesets). Previously, in Lasser et al., we used the intersection of Fu et al. and Bekker-Jensen et al. for all geneset enrichment tests, which explains why the results for the O’Neill et al overlaps are slightly different. However, the results remain significant with either background (Gheiratmand et al. 2019).

We further evaluated the significance of enrichment of autism-associated genesets in cytoskeleton-associated genesets through permutation testing. We generated 200,0000 permuted genesets for each autism-associated geneset as follows:

● “Autism genes”: we generated 200,000 permuted sets of n=255 unique genes from all genes assessed in Fu et al. 2022 (Fu et al. 2022). We weighted likelihood of gene selection by the magnitude of genetic association with autism. Specifically, we ranked genes by their magnitude of genetic association with autism as reported in Fu et al. 2022 (Supplementary Table 11, column “FDR_TADA_ASD”, (Fu et al. 2022)), where higher rank reflected greater genetic association with autism.
● “Autism PPI (baits and prey)”: We generated 200,000 permuted sets of n=100 bait and n=1043 prey. For the sets of n=100 bait, we selected n=100 unique genes from all genes assessed in Fu et al. 2022 (Fu et al. 2022) and weighted likelihood of gene selection by the magnitude of genetic association with autism as above. For the sets of n=1043 prey, we generated sets pf n=1043 random prey that matched observed prey in protein size and for which odds of selection was weighed by HEK293T protein expression levels. We defined HEK293T protein expression level by average iBAQ expression (Bekker-Jensen et al. 2017 Supplementary table 7, average of iBAQ scores for the two HEK293T experimental replicates (Bekker-Jensen et al. 2017)) and used cDNA length reported in Satterstrom 2020 as a proxy for protein size (Satterstrom et al. 2020 Supplementary Table 2, “Autosomal” sheet, “cDNA” column;). We divided the ∼10,000 HEK293T expressed proteins into 5 groups based on cDNA length (cDNA_quantile). For each observed prey, we selected a random prey from HEK293T-expressed proteins in the same cDNA quantile, with probability of selection weighted by HEK293T protein expression level.
● “Autism PPI (prey only)”: We generated 200,000 permuted sets of n=1043 prey as described above.

We evaluated whether each of the permuted autism-related genesets are significantly enriched in each of the four cytoskeleton-associated genesets using Fisher exact tests with the background gene universes described above. We calculated permuted p-values by dividing the number of permuted p-values that were as or more significant than the observed p-values by 200,000. We corrected permuted p values for multiple hypothesis testing (Bonferroni, p.adj = permuted pval * # genesets = permuted pval * 16).

### Expression correlation of autism risk genes versus ciliated cell markers across cell types

#### Tabula Sapiens (human cell atlas) dataset

We downloaded the Tabula Sapiens 2022 (Tabula Sapiens Consortium* et al. 2022) epithelial subset single-cell RNAseq Seurat object from cellxgene (https://cellxgene.cziscience.com/e/97a17473-e2b1-4f31-a544-44a60773e2dd.cxg/) (Megill et al. 2021a) in June 2023. This dataset contains scRNAseq data for 104,148 cells across 18 tissues and 62 cell types. Gene expression levels (UMI) were previously quantile normalized. We grouped cells by “cell_type”, and used percent expression (pct.exp, percent of cells in each cell type that have detected expression of the gene of interest) as a proxy for gene expression level. We categorized cell types into specific tissues of origin based on the “tissue_in_publication” that contributes the largest proportion of cells to each cell type. We restricted our analysis to the 8 tissues (with 37 associated cell types) that had at least one cell type with above 20% expression of *FOXJ1* or *ARL13B*.

#### Siletti et al 2023 (adult brain tissue) dataset

We downloaded single cell RNAseq gene expression data from Siletti et al. 2023 (Siletti et al. 2023) for n=255 hcASD, cilia markers FOXJ1 and SYNGAP1, and mitotic markers (MKI67, TOP2A, ASPM) using the CZ CELLxGENE Gene Expression webtool (https://cellxgene.cziscience.com/gene-expression) (Megill et al. 2021b). This dataset contains scRNAseq data for ∼3 million cells and 2 tissues (brain, spinal cord; 13 cell types each). We restricted our analyses to the 13 cell types from brain tissue. We used percent expression (pct.exp, percent of cells in each cell type that have detected expression of the gene of interest) as a proxy for gene expression level.

#### Autism genes (hcASD255)

We defined a set 255 high-confidence autism genes (hcASD25) that have FDR<0.1 in the most recent ASD rare-variant based whole-exome sequencing omnibus study (Fu et al. 2022).

#### Expression correlation analysis

We calculated the Pearson correlation between pct.exp of individual hcASD255 genes and the cilia markers *FOXJ1* and *ARL13B* across the 62 Tabula Sapiens cell types. We focused on Pearson correlations with *FOXJ1* given *FOXJ1* is a more specific marker for ciliated cells and because Pearson (but not Spearman) correlations identified significant correlation between the two cilia markers *FOXJ1* and *ARL13B* (positive control). We use mitotic markers (*MKI67*, *TOP2A*, *ASPM*) as negative controls. Uncorrected p-values are presented for the analysis of *SYNGAP1* specifically, while corrected p-values are presented in the larger analysis in Figure 5.

### GFP plasmids for localization screening

All human GFP-tagged autism gene and variant plasmids were constructed by cloning cDNAs from (Wang et al. 2024) into plasmid pcDNA3.1+eGFP, maintaining the same terminal side tagged. *H2B-GFP* is in a pCS2 backbone made by Reinhardt Koester, a kind gift from Richard Harland (UC Berkeley). The only autism gene within the 30 strongest-associated that we were unable to clone due to its large size was *NF1*, so it is excluded here.

### *Xenopus* husbandry

Both *Xenopus tropicalis* and *Xenopus laevis* animals were used in this study in accordance with approved UCSF IACUC protocol AN199587-00A. Ovulation was induced with human chorionic gonadotropin (Sigma CG10), and embryos were collected and manipulated according to standard procedure (Sive, Grainger, and Harland 2000). Localization experiments were done in *Xenopus laevis*, while knockdown experiments were done in *Xenopus tropicalis*. Xenbase (RRID:SCR_003280) was used for anatomical resources, phenotypes, and genetic references (M. Fisher et al. 2023; M. E. Fisher et al. 2022; Segerdell et al. 2013). Wildtype frogs were supplied by the National Xenopus Resource Center (RRID: SCR_013731) (Pearl et al. 2012).

### Human GFP-tagged protein localizations in *Xenopus laevis*

Plasmids were injected individually into 1 blastomere at the 4-cell stage along with 100 - 500 pg of Centrin-CFP (T. J. Park et al. 2008; Antoniades, Stylianou, and Skourides 2014)(Kerr, Bhan, and Héon 2016; Ozonoff et al. 1999)(T. J. Park et al. 2008; Antoniades, Stylianou, and Skourides 2014) mRNA. Several concentrations of plasmid were tested for each plasmid, ranging from 10 - 50 pg per embryo. Some plasmids were less brightly expressed than others, so the amount injected was varied to the lowest amount visible without causing overexpression ciliary phenotypes. Plasmids were also injected alone to ensure that the observed localizations were not due to the presence of Centrin-CFP or due to potential spectral overlap. For *SYNGAP1* patient variant localizations, a maximum of 20 pg total of plasmid was injected into 1 blastomere at the 4-cell stage. For the mCherry mixture experiments, 10 pg of each plasmid was co-injected for a total of 20 pg injected. The mixture experiments were done in the absence of any additional staining or markers to ensure no spectral overlap, and the mCherry plasmids were injected alone and did not show any signal in the GFP channel when imaged identically to the mixture experiments. The only autism gene within the 30 strongest-associated that we were unable to clone due to its large size was *NF1*, so it is excluded here.

### Endogenous localizations in *Xenopus*

Standard immunofluorescence staining protocols were used (H. R. Willsey et al. 2018c), with the exception of leaving out the bleaching step. Phalloidin-647 (Life Technologies A22287) was used at 1:250 with a 2 hour room temperature incubation with secondaries. Primary and secondary antibody information is included below.

### Live imaging in *Xenopus laevis*

Plasmids containing GFP-tagged hSTXBP1, hSYNGAP1, and hKM5B were injected individually into 1 blastomere at the 4-cell stage at 11-50 pg per embryo along with 100 - 500 pg of Centrin-CFP (T. J. Park et al. 2008; Antoniades, Stylianou, and Skourides 2014)(Kerr, Bhan, and Héon 2016; Ozonoff et al. 1999)(T. J. Park et al. 2008; Antoniades, Stylianou, and Skourides 2014) mRNA. Embryos were mounted in 0.02% tricaine (Fisher NC0872873) in PBS and imaged live using Fast Airyscan on a Zeiss LSM 980 microscope.

### *Xenopus tropicalis* whole mount in situ hybridization

Hybridized chain reaction (HCR) on *X. tropicalis* NF stage 25 embryos was performed according to (H. R. Willsey 2021) addition of heating probes to 95°C for 90 seconds and cooling to room temperature before use. *syngap1* and *foxj1* probes were custom ordered through Molecular Instruments and designed to target the *X. tropicalis* genes.

### Rat husbandry for isolation of rat striatal neurons

Timed-pregnant female Sprague-Dawley rats were obtained commercially (Charles River) and housed until embryonic day 18 (E18). Animals were housed individually on a 12-h light/dark schedule with constant access to enrichment, food and water. All experimental procedures were performed in accordance with the guidelines of the UCSF Institutional Animal Care and Use Committee (protocol number AN185688), and according to the National Institutes of Health Guide for Care and Use of Laboratory Animals.

### Isolation of rat striatal neurons

To isolate striatal neurons, a timed-pregnant female rat was euthanized (CO_2_ and bilateral thoracotomy) and embryos of both sexes were collected into ice-cold HBSS calcium/magnesium/phenol red-free (Thermo Fisher). Brains were extracted from the skulls into a fresh dish of ice-cold HBSS calcium/magnesium/phenol red-free (Thermo Fisher). The striatum (including dorsal and ventral striatum) was dissected under a stereomicroscope (Nikon) into a 15 mL falcon tube filled with ice-cold HBSS resting on ice. The tissue was dissociated in 0.05% trypsin/EDTA (UCSF Media Production) for 15 min at 37°C with periodic agitation. The tissue was washed 2x in DMEM (Thermo Fisher) supplemented with 10% fetal bovine serum (UCSF Media Production) and 30 mM HEPES (Gibco). Neurons were further dissociated by trituration through 2 different sterile pipettes with sequentially smaller diameter openings. 200,000 neurons were seeded onto a well in a 24 well glass bottom plate with high performance #1.5 coverglass (Cellvis) coated with poly-D-lysine (Sigma). On DIV 4-5 medium was exchanged for phenol red-free Neurobasal medium (Thermo Fisher) supplemented with GlutaMAX 1x (Thermo Fisher) and B-27 1x (Thermo Fisher). Half of the culture medium was exchanged every week with fresh, equilibrated medium. On DIV 8-9, neurons were transfected with Lipofectamine 2000 (Thermo Fisher). For each transfection reaction 2 μl of Lipofectamine was combined with 1–2 μg DNA in 0.5 ml of media per well of a 24 well plate, and media was exchanged 4–6 h later. Cytosine arabinosine 2 μM (Millipore Sigma) was added at DIV 8-9 to reduce growth of non-neuronal cells. Neurons were maintained in a humidified incubator with 5% CO_2_ at 37°C. All experiments were performed on at least 3 independent cultures.

### Human GFP-tagged protein localizations in rat striatal neurons

To generate Flag-tagged DRD1 receptor plasmid, the pCAGG-G418 vector (addgene #31264, A. Pierce Lab) was modified by inserting a signal sequence (SS) and FLAG sequence at EcoRI and NotI sites. The mouse dopamine receptor DRD1 was purchased as a gene block from IDT and was inserted into the pCAGG-G418 SS Flag vector at the NotI site as previously reported (Fagan et al. 2024). To label neuronal primary cilia, transfected Flag DRD1 receptors were surface labeled with a rabbit anti-Flag antibody (Millipore Sigma) at 37°C for 15 minutes. Briefly, the FLAG antibody was diluted 1:1000 in conditioned Neurobasal media. Immediately following, neurons were fixed in 4% paraformaldehyde and 4% sucrose for 10 min at room temperature and then washed with PBS-T (0.2% Triton X-100 in PBS). Neurons were blocked in 3% normal goat serum (Millipore Sigma) and 0.3% Triton X-100 diluted in PBS at RT for 1 hr. Primary antibodies (see below) were diluted into blocking solution and incubated overnight at 4°C. The following day, neurons were washed 3 times in PBS-T. Secondary antibodies were diluted in blocking solution (see below) and incubated for 1 hour at RT. Neurons were washed in PBS-T. Nuclei were stained with Hoechst 33342 (Thermo Fisher) at 1:10,000 diluted in PBS for 10 minutes at RT. Following, neurons were washed 2 times in PBS and stored in 1 mL PBS at 4°C until imaging. The only autism gene within the 30 strongest-associated that we were unable to clone due to its large size was *NF1*, so it is excluded here.

### Endogenous protein localizations in rat striatal neurons

Neurons were fixed in 4% paraformaldehyde and 4% sucrose for 10 min at room temperature and then washed with PBS and stored in PBS at 4°C until staining and imaging. Neurons were permeabilized in for 30 minutes in PBS-T (0.2% Triton X-100), 3x for 5 minutes, and blocked for at least 30 minutes in 3% BSA (VWR 0332) or 10% normal donkey serum (Southern Biotech 0030-01) diluted in 0.2% PBS-T. Primary antibodies (see below) were diluted into blocking buffer and incubated for 3 hours at room temperature with gentle agitation. Secondary antibodies (see below) were diluted in blocking solution at 1:1000 and incubated for 1 hour at room temperature. Neurons were washed 3X for 5 minutes in PBST. ARL13B-647 (1:100, CL647-17711 (Proteintech)) was diluted 1:500 in blocking buffer and incubated overnight at 4°C. Neurons were washed once with PBST and nuclei were stained with DAPI (Thermo Fisher Scientific D3571) at 1:1000 diluted in PBS for 30-45 minutes at room temperature. Following neurons were washed 3X for 5 mins in PBS and stored in 0.5 mL of PBS 4°C until imaging. Neurons were seeded on 24-well glass bottom dishes with no. 1.5 coverslips coated with Fibronectin (Invitrogen) or poly-d-lysine (Cultrex). The medium was changed to one without phenol red.

### Endogenous localizations in rat brain ependymal cell cilia

Long Evans outbred rats were maintained in the Kastner laboratory under UCSF IACUC-approved protocol AN193303. Adult male rats (4 months old) were used. Animals were euthanized and perfused, after which the brain was dissected and post-fixed in 4% paraformaldehyde for 24 hours, followed by equilibration to 30% sucrose solution over 3 days at 4°C. Brains were then cryopreserved and sectioned into 10 µm sections using a Thermo Scientific CRYOSTAR NX50 cryostat. Sections containing the lateral ventricles were stained on slides by standard immunofluorescence protocol with permeabilization in 0.2% Triton-X100 detergent in PBS (PBT), 1 hour blocking in 2% BSA (VWR Life Science 0332) in PBT, overnight primary antibody in blocking solution, and 2 hour room temperature secondary antibody in blocking solution incubation. Primary and secondary antibodies used are listed below.

### *Xenopus tropicalis* gene perturbations and rescues

For *Xenopus tropicalis* knockdown experiments, CRISPR sgRNAs were synthesized with a EnGen kit (NEB E3322S) and purified (Zymo R1018), or morpholinos were purchased from Gene Tools. Morpholino sequences are: *chd8* (5’-CCAGCCTGTGAGAGAAGATAGTAAT-3’), *chd2* (5’-GGTTTATCCTCATTCCTCATCATTG-3’), *pogz* (5’-TTCCATAAACAGATCCGTGTCCATG-3’), *syngap1* (5’-GAGCATAGAACATCATTCCACAGCT-3’), and standard control (5’-CCTCTTACCTCAGTTACAATTTATA-3’). *syngap1* sgRNA sequence is (5’-TTCTAATACGACTCACTATAGCAGAGGGTGGACAACGCATGTTTTAGAGCTAGA-3’). For the chromatin regulator morpholinos (*chd8*, *chd2*, *pogz*), 0.4 pmol of morpholino was injected into 1 blastomere of 2-cell stage *Xenopus tropicalis* embryos, along with 50 pg of tracer Centrin-CFP mRNA. For *syngap1* morpholino, 1 pmol of morpholino was injected with 300 pg of tracer Centrin-CFP mRNA into 1 blastomere of 4-cell stage *X. tropicalis* embryos. For *syngap1* morpholino rescue, 8.6 pg of hSYNGAP1-GFP plasmid was co-injected with the same amount of morpholino and Centrin-CFP tracer mRNA. For *syngap1* CRISPR, 5 ng Cas9-NLS protein (UC Berkeley Macrolabs) was injected with 2.8 ng sgRNA and 1 ng of tracer Centrin-CFP mRNA into 1 blastomere at the 2-cell stage in *X. tropicalis*. Crispant embryos were incubated for 1 hour at 28°C and then overnight at 25°C.

### *Xenopus* ciliary phenotyping

For chromatin regulators, positively-injected ciliated cells were identified by positive Centrin-CFP signal and the acetylated-tubulin staining was categorized according to phenotype. At least 3 representative embryos per condition were assessed, with 20 cells total per condition. For *syngap1*, positively-injected embryos were identified by positive Centrin-CFP signal and embryos themselves were categorized as having normal ciliation or having cilia defects affected based on their acetylated tubulin staining. For the morpholino rescue experiment, at least 20 representative embryos per condition were assessed. Cilia were assessed between stage 27 and stage 35. We conducted Fisher’s Exact Tests (two-sided) using a 3×2 contingency table using the injection status (uninjected, *syngap1* morpholino, *syngap1* morpholino with the hSYNGAP1-GFP) as rows and the ciliary phenotype (no ciliary phenotype vs. severe ciliary phenotype) as columns (Table S5). We then performed pairwise comparisons using Fisher’s Exact test and we corrected p values for multiple hypothesis testing (Bonferroni). Prism software (v10.1.1) was used to graph and R was used to determine statistical significance between conditions.

### iPSC-derived neurons

Lentivirus was produced using LentiX-293T cells (Clontech) that were maintained in DMEM with Glutamax (Fisher Scientific 10566016), MEM-NEAA (Fisher Scientific 11140-050) and 10% FBS. Lentiviral packaging was performed by seeding 400,000 cells per well of a 6-well plate, then transfecting with 1 µg equimolar packaging mix (pMDL, Addgene 1251; pRSV, Addgene1253; pVSV-g) and rtTA (phr_tet3g) or pLV-TetO-hNGN2-eGFP-Puro using OptiMEM and Lipofectamine 2000 (Fisher Scientific 11668019). On the day of transduction (48 hours later), viral supernatant was harvested from Lenti293T cells, filtered through a syringe-driven 0.45µM filter (Millex-HP, Cat. No. SLHPR33RS) into 15mL Falcon tubes.One volume of Lenti-X concentrator (Takara Bio, 631231) was combined with three volumes of clarified supernatant, incubated for 30 minutes, centrifuged at 1,500 g for 45 minutes at 4 degrees Celsius and resuspended in appropriate amount of mTESR Plus media containing Rock inhibitor (0.5 mL) One volume of Lenti-X concentrator (Takara Bio, 631231) was combined with three volumes of clarified supernatant, incubated for 30 minutes, centrifuged at 1,500 g for 45 minutes at 4 degrees Celsius and resuspended in appropriate amount of mTESR Plus media containing Rock inhibitor (0.5 mL).

Deidentified human iPSCs were obtained from Rarebase for a female unaffected control line (RB-FM45-PT1), which has a typical karyotype. iPSCs were cultured in mTESR Plus media (with supplement, Stem Cell Technologies 100-0276) on Corning dishes (6-well dishes Cat. No. 3516 and 10 cm dishes Cat. No. 430293 for expansion) coated with Corning Matrigel hESC-Qualified Matrix (Corning 354277) diluted in mTESR Plus media. Briefly, mTESR Plus media was replaced every day. When 80-90% confluent, cells were passaged as follows: Media was aspirated, cells were washed twice with PBS (Fisher Scientific 10-010-049), 1 mL ReLeSR (Stem Cell Technologies, 100-0483) per well of a 6-well plate was added, incubated at room temperature for 30 seconds, ReLeSR was aspirated off, cells were incubated at 37°C for 7 minutes; 3 mL of mTESR media was added; the detached cell aggregates were collected,transferred to a 15 ml conical tube, resuspended cell multiple times to obtain small cell aggregates and passaged at a required ratio.

iPSCs were differentiated into cortical excitatory neurons using NGN2-induction. iPSCs were lentivirally transduced with pLV-TetO-hNGN2-eGFP-Puro (Addgene 79823), which contains doxycycline-inducible human NGN2, enhanced GFP, a puromycin resistance gene, and a tet-transactivator (phr_tet3g). On the day of transduction, iPSCs were passaged into single cells using accutase as follows: Washed cells twice with 1X PBS (ph 7.4), incubated with 1 mL accutase per well of a 6-well plate for five minutes, add 1 mL PBS and collect cells into a 15 mL conical tube; collect residual cells with another 2 mL PBS. Cells were counted and resuspended in virus-containing media (described above) at a ratio of 1 million iPSCs to 1 mL of total virus-containing supernatant (0.5 mL of pLV-TetO-hNGN2-eGFP-Puro and 0.5 mL phr_tet3g supernatant). Cells were expanded and passaged twice using ReLeSR as described above. At passage 11, NGN2 and puromycin resistance expression was induced as follows: Seven million cells were passaged into pre-differentiation media containing Rock inhibitor (Y-27632, Dihydrochloride, Stem Cell Technologies 72304) and doxycycline (Sigma D3447) as described in Tian et al (Tian et al. 2019; Kampmann 02 2020) into a 10 cM dish using accutase as described above. Media was replaced completely daily (Day −3, Day −2, Day −1) with media containing doxycycline. On Day 0, cells, 70,000 cells per well were passaged into N2/B27 selection media (Tian et al. 2019; Kampmann 02 2020) containing doxycycline and puromycin using accutase onto glass bottom 24-well plates with high performance #1.5 coverglass (Cellvis P24-1.5H-N) coated with Poly-L-Ornithine diluted (Sigma P4957, 15 ug/mL, diluted in culture grade water) at least 24 hours in advance and laminin (Gibco 23017-015, final concentration 1 ug/mL diluted in 1X PBS) and fibronectin (Sigma F1141, final concentration 2 ug/mL diluted in 1X PBS) at least 3 hours in advance. Cells underwent a full media change with puromycin in the media on Day 3 and cells were fixed with 4% paraformaldehyde (final concentration) on Day 7, washed with PBS three times, and stored at 4°C for further processing.

### Endogenous localization in iPSCs

iPSCs (cultured and fixed as above) were permeabilized with PBST (1X PBS (Fisher Scientific, 10-010-049) at pH 7.4 with 0.2% Triton X-100) 3 times for 15 minutes. Cells were blocked with 10% CAS-Block (Life Technologies 8120) in PBST for 2 hours at room temperature. Primary antibodies (see below) were diluted into this blocking solution, and cells were incubated at 4°C for 36 hours. Following primary antibody incubation, cells were washed 3 times for 15 minutes in PBST and 2 times in PBS, protected from light. Fluorophore conjugated secondary antibodies and stains (see below) were diluted in blocking solution and cells were incubated for 1 hour at room temperature, protected from light. Cells were washed with PBST 3 times for 15 minutes and three times with PBS for 5 minutes; cells were stored at 4°C, protected from light.

### Antibodies and Stains

Primary antibodies used: acetylated alpha-Tubulin (Sigma T6793, 1:600 for *Xenopus*, 1:1000 for rat brain slices), ARL13B (BiCell Scientific 90413h, 1:100 for human iPSC-derived neurons), CHD8 (Abcam ab114126, 1:100 for *Xenopus* and for rat striatal neurons), CHD2 (Novus Biologicals NBP2-32563, 1:100 for *Xenopus* and for rat striatal neurons), POGZ (Abcam ab167408, 1:100 for *Xenopus* and for rat striatal neurons), SYNGAP1 (MyBioSource MBS3216405, 1:100 for *Xenopus*), SYNGAP1 (Abcam ab3344, 1:1000 for rat ependymal cells or human iPSC-derived neurons), SYNGAP1 (Invitrogen PA1-046, 1:500 for rat striatal neurons), STXBP1 (Synaptic Systems 116003, 1:500 for rat striatal neurons, 1:150 for rat ependymal cells), SHANK3 (Santa Cruz Biotechnology, sc-377088, 1:1000 for rat ependymal cells), PSD95 (Santa Cruz Biotechnology, sc-32290, 1:200 for rat striatal neurons, 10% donkey serum block), SCN2A (Alomone labs ASC-002, 1:100 for rat striatal neurons), SLC6A1 (Novus Biologicals NBP1-89802, 1:100 for rat striatal neurons), GRIN2B (Abcam ab65783, 1:200 for rat striatal neurons), FGFR1OP (Abnova H00011116-M01, 1:500), FGFR1OP (ProteinTech 11343-1-AP, 1:100), ARL13B-647 (ProteinTech CL647-17711, 1:500), and anti-GFP (Novus NB100-1614, 1:1000).

Secondary antibodies include: anti-mouse AlexaFluor-488 (Thermo Fisher A32723), anti-mouse AlexaFluor-555 (Thermo Fisher A32727), anti-mouse AlexaFluor-647 (Thermo Fisher A32728), anti-rabbit AlexaFluor-488 (Thermo Fisher A32731), anti-rabbit AlexaFluor-555 (Thermo Fisher A32732), anti-rabbit Alexa Fluor 594 (Thermo Fisher A11012), anti-rabbit AlexaFluor-647 (Thermo Fisher A32733), anti-rat AlexaFluor-488 (Thermo Fisher A48269), anti-chicken Alexa Fluor-488 (Thermo Fisher A32931). Secondary antibodies were used at: 1:250 in *Xenopus*, 1:1000 in rat tissue, 1:2000 in rat cultured neurons, and 1:1000 in human iPSC-derived neurons.

Phalloidin-647 (1:250, Life Technologies A22287) and/or DAPI (1:1000, Thermo Fisher Scientific D3571) were incubated with secondaries.

### Microscopy

All images presented in the figures were acquired on a Zeiss LSM 980 confocal microscope with Plan Apo 63x oil objective, 1.6 numerical aperture (NA) oil objective, and a 405-nm, 488-nm, 555-nm, and/or 647-nm laser with the exception of Figure S7 which was acquired on a Leica SP8 confocal microscope with a 63X oil objective. Images were acquired either in confocal mode or with Zeiss Airyscan processing. Images were processed in Zen or ImageJ. Bandpass filters were utilized in confocal mode to minimize spectral overlap. For making a determination on whether human GFP-tagged autism proteins localize to neuronal rat cilia, cells were imaged as above or by confocal microscopy using a Nikon Ti inverted fluorescence microscope with CSU-22 spinning disk confocal (Yokogawa) with a PlanApo VC 60X 1.4 NA oil objective. For images presented in rat striatal neuron localization screen (Fig. 2-3), upper and lower limits for 488 laser power were determined by imaging N and C terminus GFP controls alone, and all images were imaged using 488 laser power within that range and presented as average intensity projections. For images presented in *Xenopus* localization screen figures (Fig. 2-3), an upper limit for 488 laser power was determined by imaging H2B-GFP controls alone, and all images were imaged using 488 laser power below that value and presented as maximum intensity projections of selected slices. Nuclear Z-planes depict a maximum intensity projection of slices at the nuclear plane. Brightness and contrast was adjusted for each whole image to be representative.

500 nm Tetraspeck beads (Thermofisher, T7281) and Zen Blue Channel Alignment (Extended) function were used to aid in channel alignment as needed. For imaging, cells were seeded on 24-well glass bottom plates with high performance #1.5 coverglass and *Xenopus* embryos were mounted in vacuum grease wells on slides with #1.5 coverslips in PBS.

### Patient medical record review

Citizen data for *SCN2A*, *STXBP1*, *CHD2*, *SYNGAP1*, and *SLC6A1* were obtained with IRB non-humans subject research determination (IRB #23-39079). Patients were included in analysis if they had a genetic variant in the gene being studied that was classified as “Pathogenic,” “Likely Pathogenic” or “Variant of Unknown Significance” in their genetic report. Clinical diagnoses were manually annotated as “cardiac,” “dysmorphic body,” “dysmorphic face,” “kidney,” “respiratory,” “seizure,” “autism,” “skeletal,” or “other.” When multiple annotations were relevant, clinical diagnoses were assigned a secondary or tertiary annotation. Clinical diagnoses were considered to be a part of a category if they had a primary or secondary annotation in that category. Diagnoses annotated as “cardiac” were further annotated as being “structural,” “arrhythmia,” “murmur,” “heart conduction disorder,” or “valve regurgitation.” All other heart diagnoses had less than 3 instances and were combined as “other.” Prevalence of structural heart defects was calculated by summing the unique number of patients with “structural” and/or “heart murmur” annotations and calculating a percentage using the total number of patients analyzed. Data analysis and visualization were performed in GraphPad Prism software.

### nNO measurements and analyses

This work was conducted under the advisement of the UCSF Institutional Review Board with approval #23-39385. The patient’s caregiver provided: nature of the patient’s genetic variant, ages, and sexes. Patient and family member (sibling or parent) values were recorded using a NIOX VERO nNO machine by tidal breathing method using adult or pediatric olives, as appropriate. Tidal breathing method, which requires minimal cooperation compared to the forced exhale method, was selected since many of these patients also have intellectual disability. Of the 26 patients for whom testing was attempted, only 2 were unable to complete the test. All unaffected family members were able to complete the test. This machine has been shown to discriminate PCD with 100% specificity (Jõgi et al. 2023), and a general cutoff of < 77 nL/min (< 257 ppb on this machine) by forced exhale for PCD diagnosis has been proposed (Shapiro et al. 2018), though many consider a lower cutoff of 30 nL/min (< 100 ppb on this machine) for the tidal breathing method (Beydon et al. 2023; Paternò et al. 2023). Whether differences in nNO were significantly different between groups was assessed by nonparametric, one-tailed, paired and unpaired Wilcoxon tests, as described. We had the hypothesis that patients would have lower nNO values going into the study, motivating the choice of a one-tailed test.

## Supplementary Materials

**The PDF file includes:**

Figures S1 to S12

Table Legends for Tables S1 to S5

**Figure S1.**
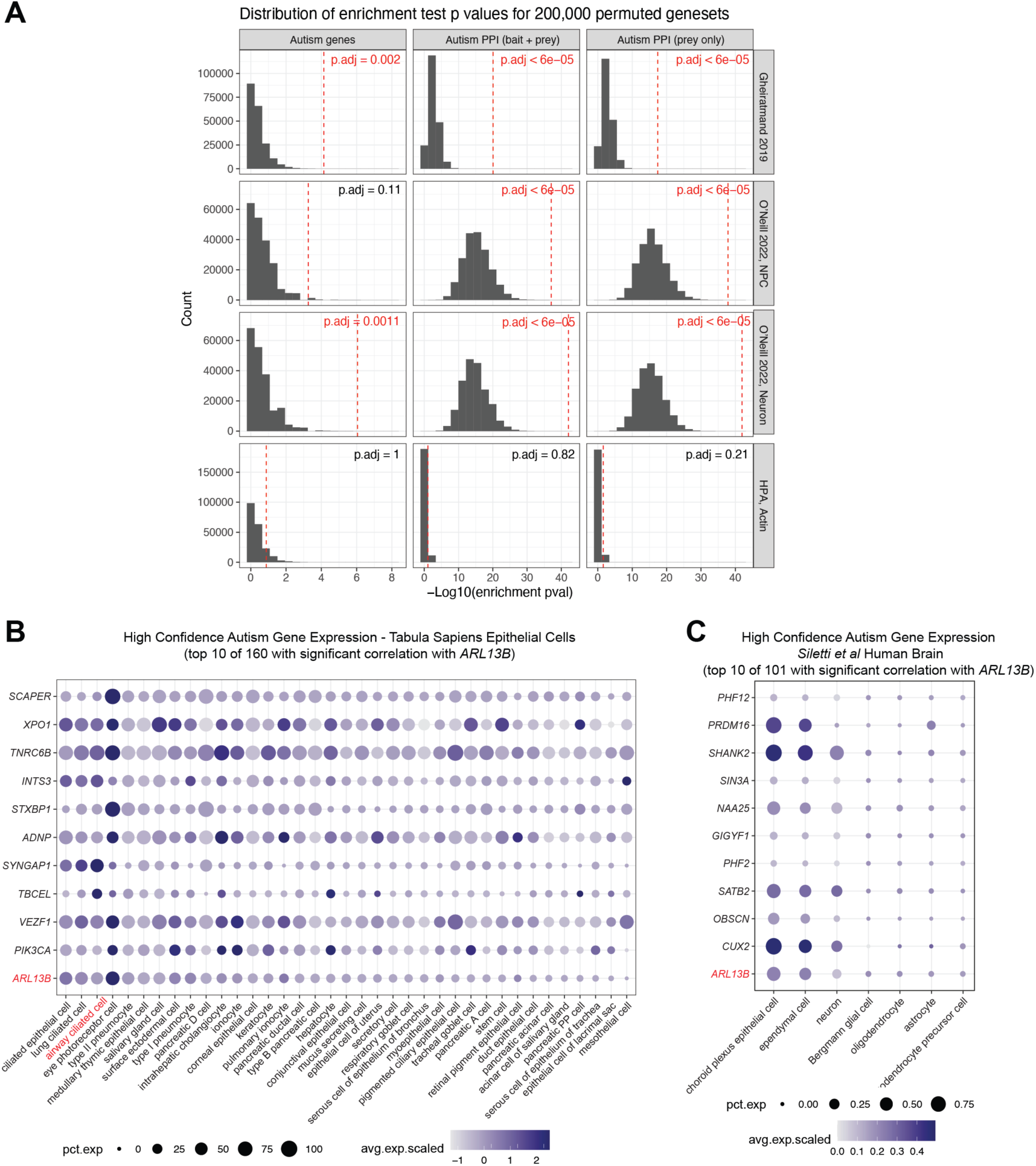
PPI network enrichment permutation testing and gene expression correlations. (A) Permutation test results for enrichments of autism proteins within cytoskeletal proteomes. (B) Quantile-normalized gene expression levels of top ten high confidence autism genes with significant correlation with ciliary marker *ARL13B* in epithelial cell types found in the *Tabula Sapiens* dataset. Expression frequency (percent of cells within a given cell type expressing the gene) is indicated by dot size and scaled expression level is indicated by color intensity. (C) Quantile-normalized gene expression levels of top ten high confidence autism genes with significant correlation with ciliary marker *ARL13B* in brain cell types found in the *Siletti et al* dataset. Expression frequency (percent of cells within a given cell type expressing the gene) is indicated by dot size and scaled expression level is indicated by color intensity.

**Figure S2.**
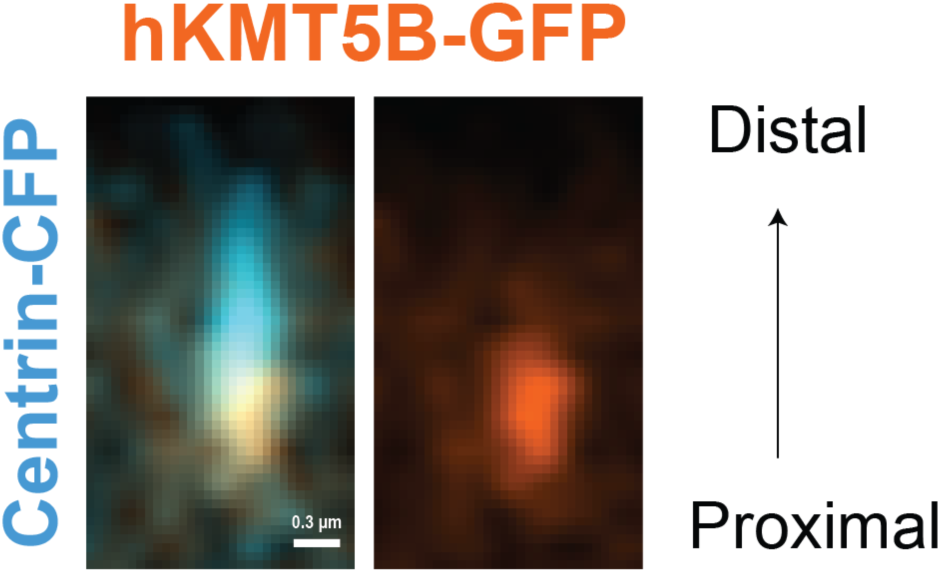
Autism-associated chromatin regulator KMT5B localizes proximal to the basal body. Human GFP-tagged KM5TB (orange) localizes proximally at the basal body (labeled by centrin-CFP, cyan), imaged live.

**Figure S3.**
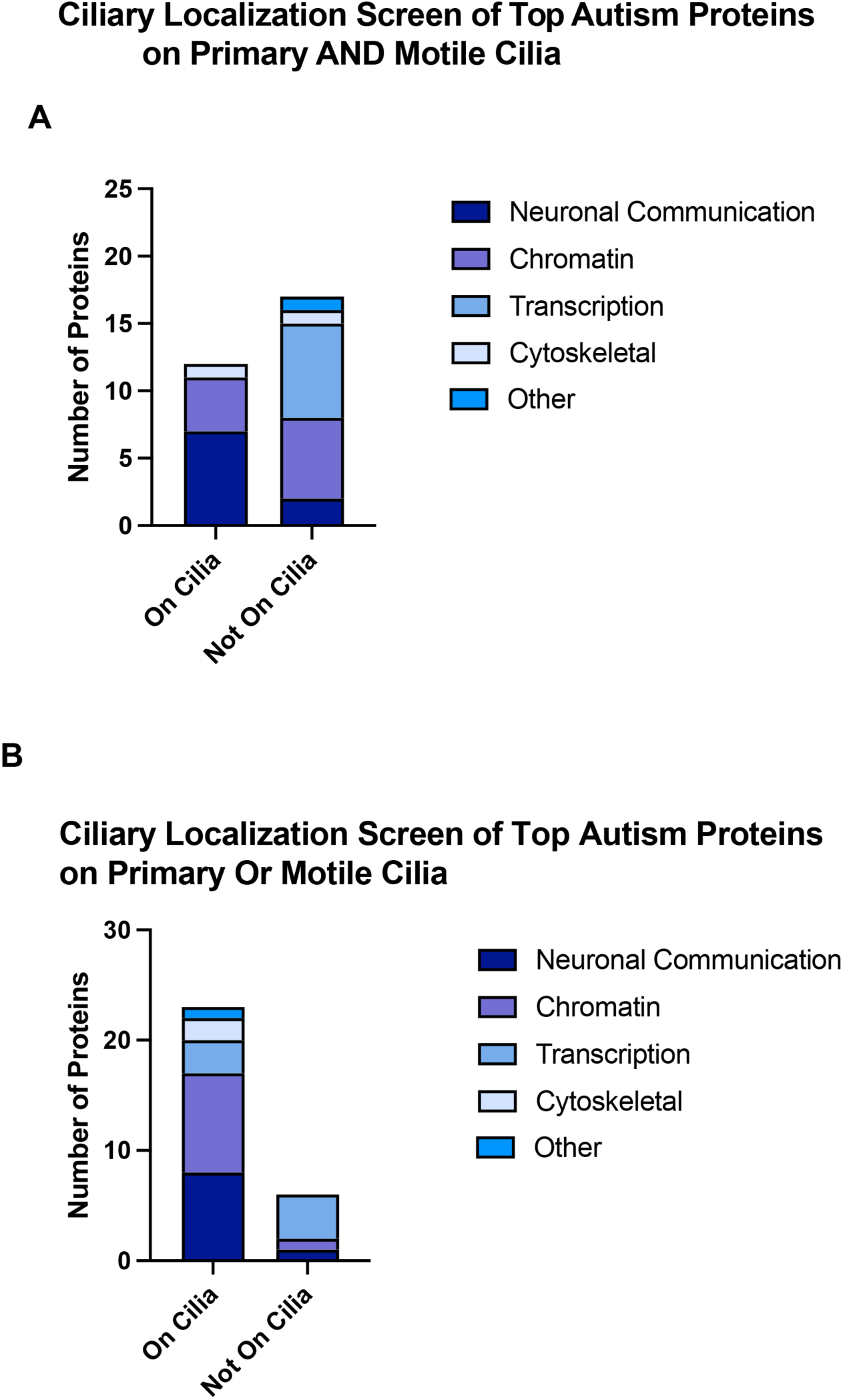
Results of ciliary localization screen of top autism proteins by functional annotation. (A) Quantification of the number of autism-associated proteins that localize to primary cilia and motile cilia, with each functional annotation depicted in a different color. (B) Quantification of the number of autism-associated proteins that localize to primary cilia or to motile cilia, with each functional annotation depicted in a different color. One gene, *NF1*, was unable to be cloned due to its large size and is excluded here.

**Figure S4.**
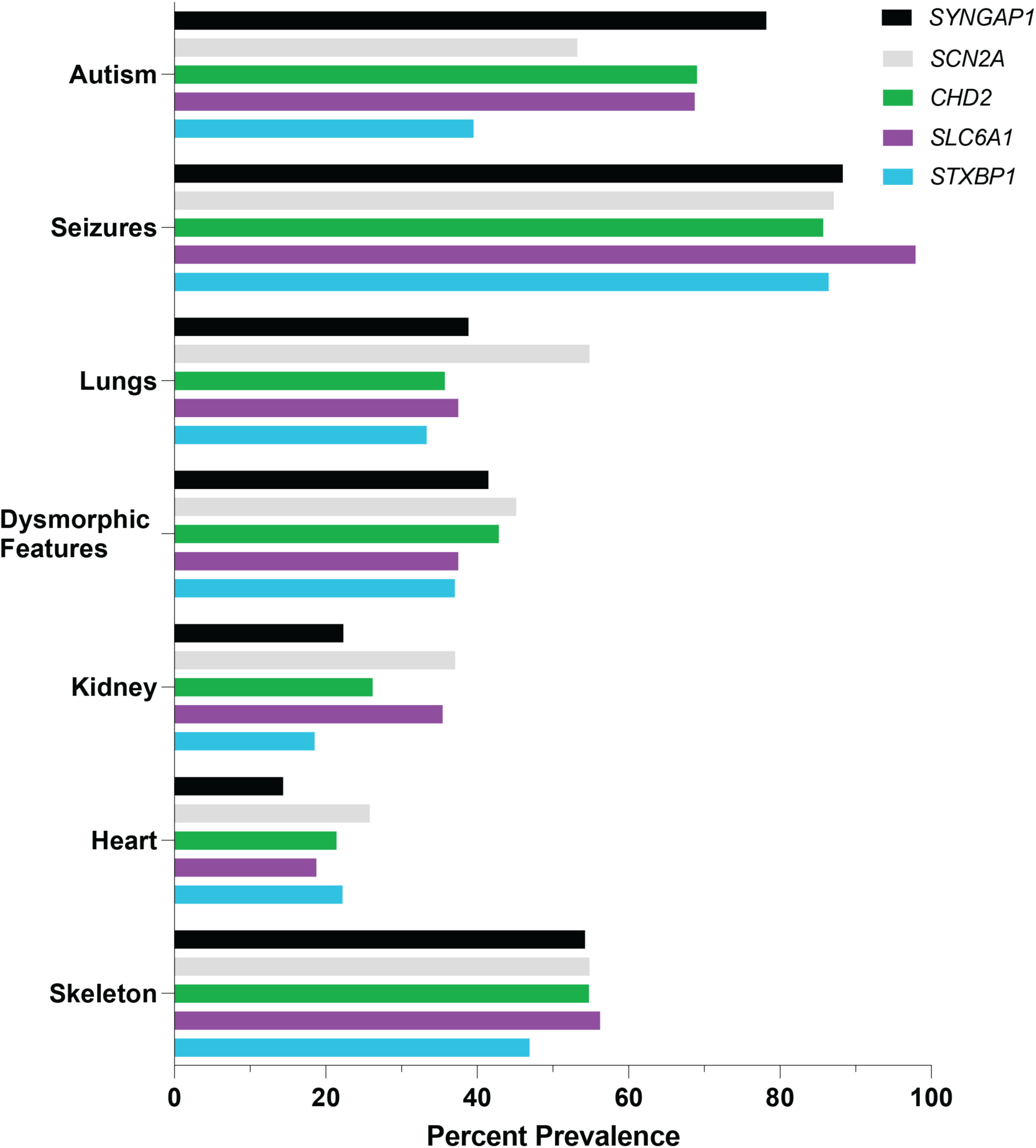
Summary-level results from Citizen patient medical record data review for 5 high confidence autism gene patient cohorts. Citizen patient medical record data shows high prevalence of autism, seizures, respiratory issues, dysmorphic features, kidney issues, cardiac issues, and skeletal defects in patients with variants in SYNGAP1, SCN2A, CHD2, SLC6A1, STXBP1 that are annotated as not benign or likely benign.

**Figure S5.**
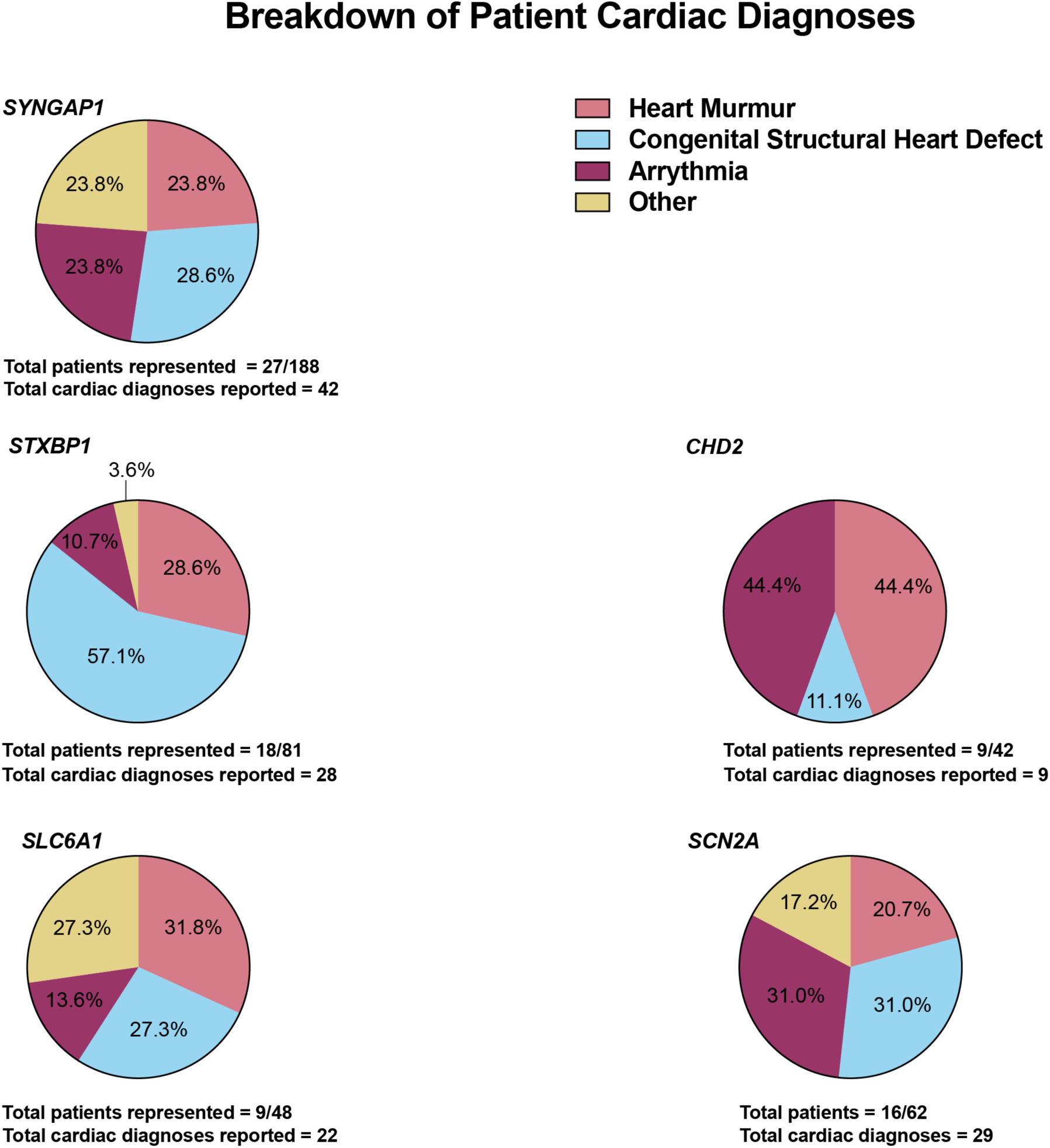
Citizen patient medical record data specifically for cardiac diagnoses. More than 50% of the cardiac phenotypes observed in patients with *de novo* variants in *SYNGAP1, STXBP1, CHD2, SLC6A1,* and *SCN2A* are structural heart defects or murmurs. Cardiac diagnoses percentages are displayed for all patients with *de novo* variants in a given gene (labeled above each chart). Total number of unique patients with cardiac diagnoses is listed below each chart; the total number of cardiac diagnoses reported for all patients with non-benign or non-likely benign variants are also shown for each chart, accounting for those with more than one observed cardiac diagnosis.

**Figure S6.**
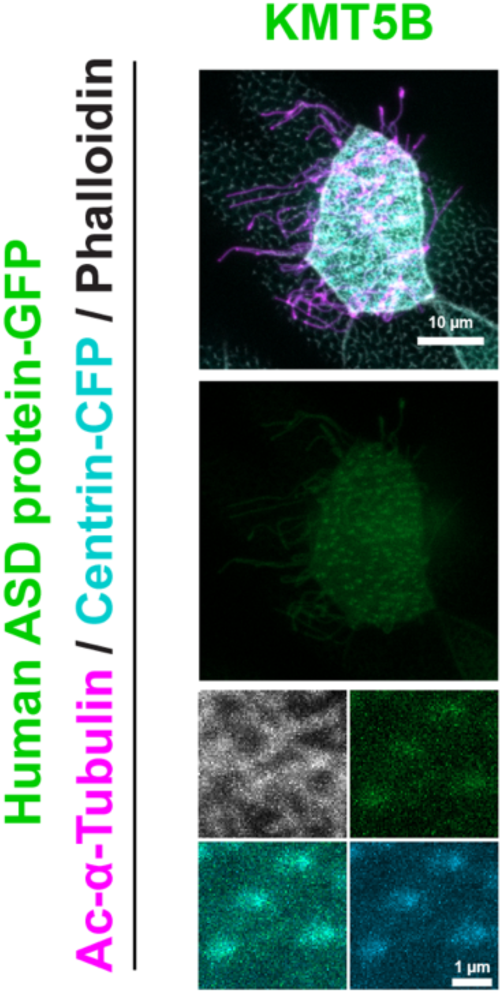
Autism-associated chromatin regulator KMT5B localizes to motile cilia. Human GFP-tagged construct for autism-associated chromatin regulators KMT5B (green) localizes to motile cilia (labeled by acetylated alpha-Tubulin, magenta; basal bodies labeled by Centrin-CFP, cyan) when expressed in the *X. laevis* embryonic epidermis.

**Figure S7.**
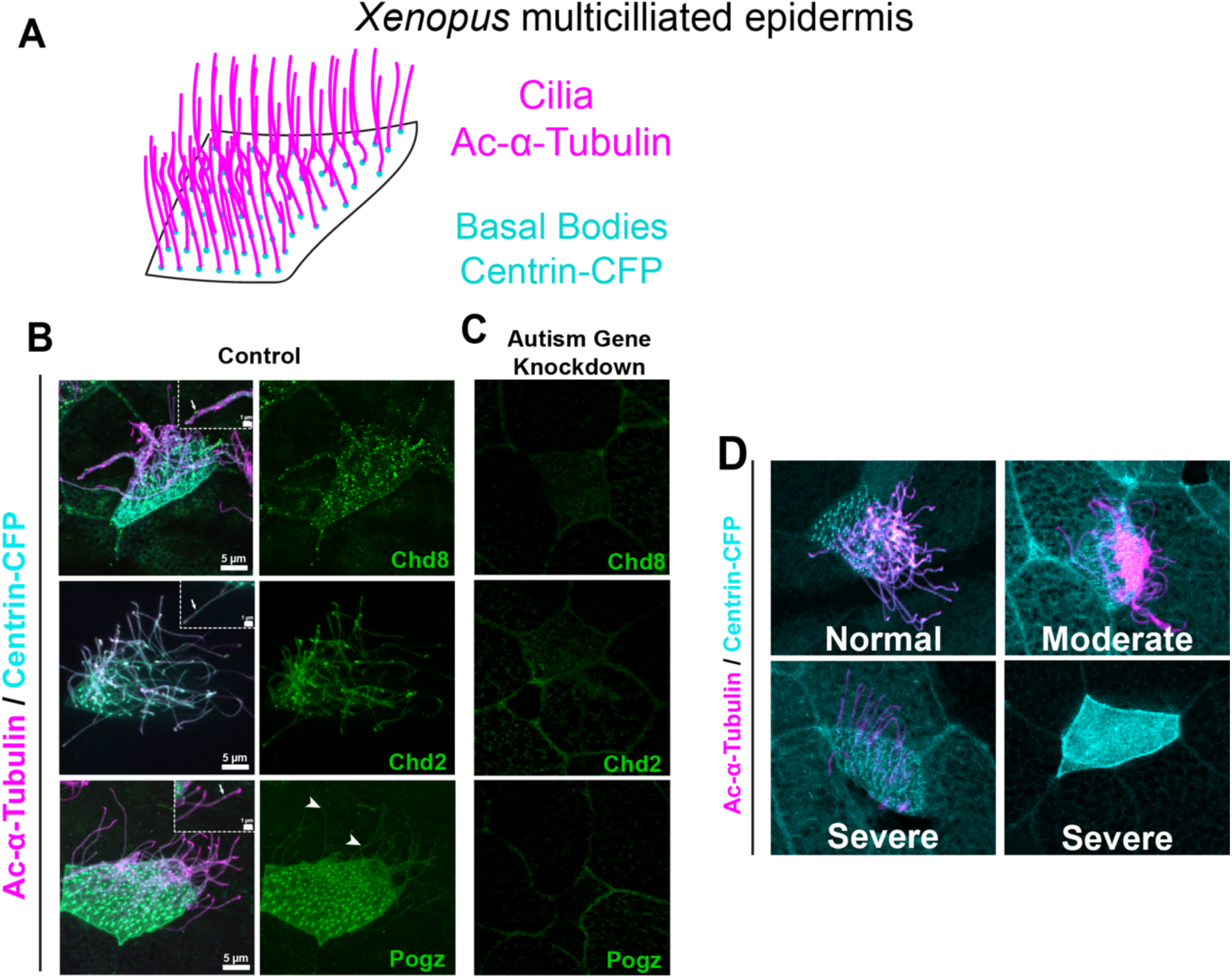
Localization and phenotyping of autism-associated chromatin regulators. (A) Schematic of *Xenopus* multiciliated epidermis with immunofluorescence labeling scheme in which ciliary axonemes are labeled by acetylated alpha tubulin in magenta and basal bodies are labeled by injecting centrin-CFP. (B) In control *X. tropicalis* multiciliated cells, endogenous proteins Chd2, Chd8, or Pogz (green) localize to cilia (labeled by acetylated alpha-Tubulin, magenta, and centrin-CFP, cyan). (C) Fluorescence signal for Chd2, Chd8, or Pogz is abrogated when *chd2*, *chd8* or *pogz* are knocked down. (D) Representative images of cilia (basal bodies labeled by centrin-CFP, cyan; axonemes labeled by acetylated alpha-Tubulin, magenta) with phenotypes categorized as “normal,” “moderate,” or “severe” for quantification presented in Fig. 2E.

**Figure S8.**
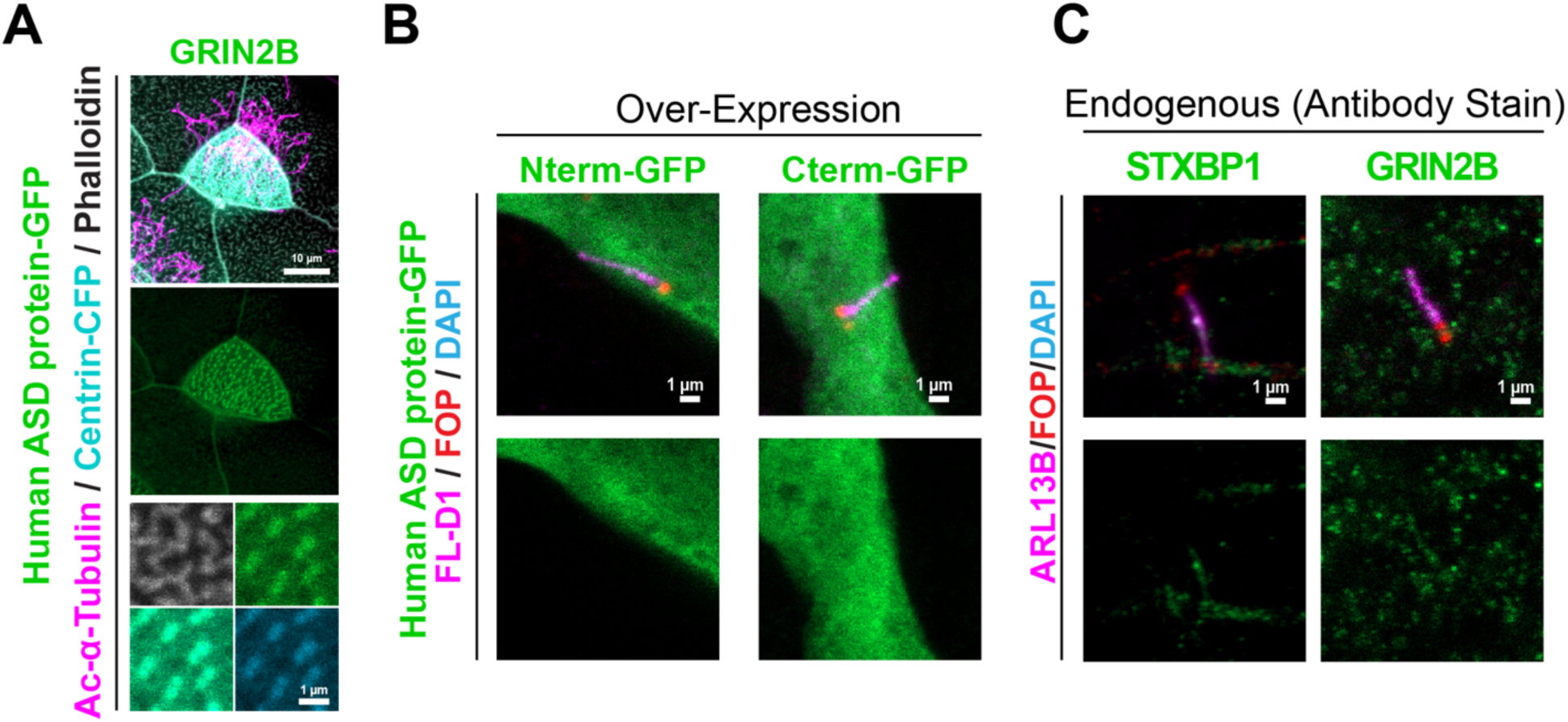
Autism neuronal communication proteins GRIN2B and STXBP1 localize to cilia, but control plasmids do not. (A) Human GFP-tagged GRIN2B protein (neuronal communication annotation) localizes to cilia (axonemes labeled by acetylated alpha tubulin, magenta; basal bodies labeled by centrin-CFP, cyan) when expressed in *X. laevis*. (B) Human GFP-tagged constructs for control plasmids containing empty vector with GFP tagged at the N or C terminus do not (green) localize to primary cilia (Flag-tagged Dopamine D1 receptor, magenta; basal bodies labeled by FGFR1OP, red) when expressed in primary rat striatal neurons. (C) Endogenous antibody staining for autism associated neuronal communication proteins STXBP1 and GRIN2B in primary cilia (labeled by ARL13B, magenta; basal bodies labeled by FGFR1OP, red) in primary rat striatal neurons.

**Figure S9.**
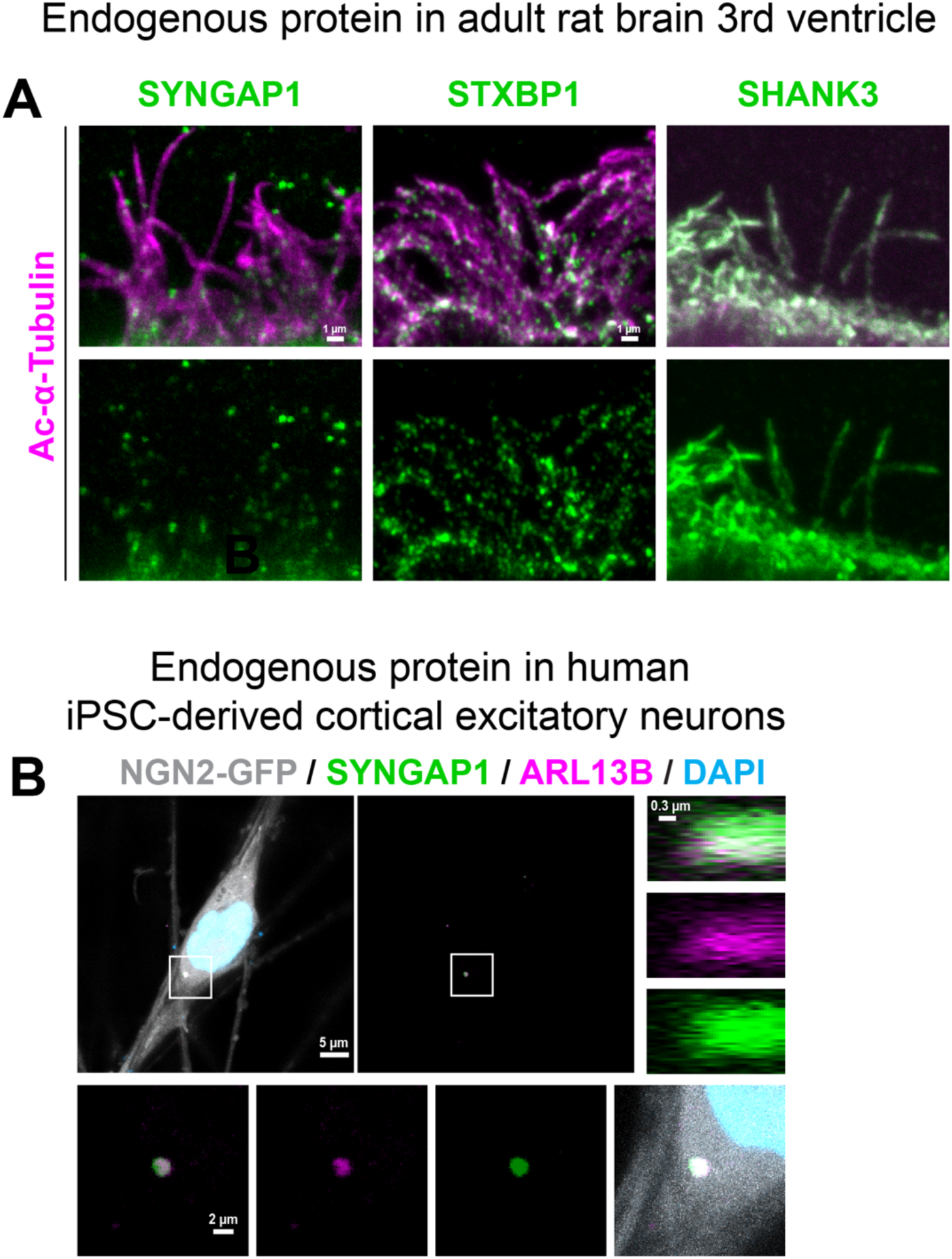
Autism-associated neuronal communication proteins localize to cilia in rat brain ependymal cells and human iPSC-derived cortical neurons. (A) Endogenous SYNGAP1, STXBP1, and SHANK3 proteins (green, labeled by published antibodies) localize to cilia (labeled by acetylated alpha-Tubulin, magenta) in ependymal cells along the rat lateral ventricles. (B) Endogenous SYNGAP1 (green, antibody staining) localizes to cilia (labeled by ARL13B antibody staining, magenta) in human iPSC-derived cortical excitatory neurons (transduced with NGN2-GFP for differentiation, gray). Nucleus is labeled by DAPI (blue). Bottom row shows cilium at higher magnification. Right panel shows SYNGAP1 (green) localizing to cilia (magenta) in the XZ plane at higher magnification.

**Figure S10.**
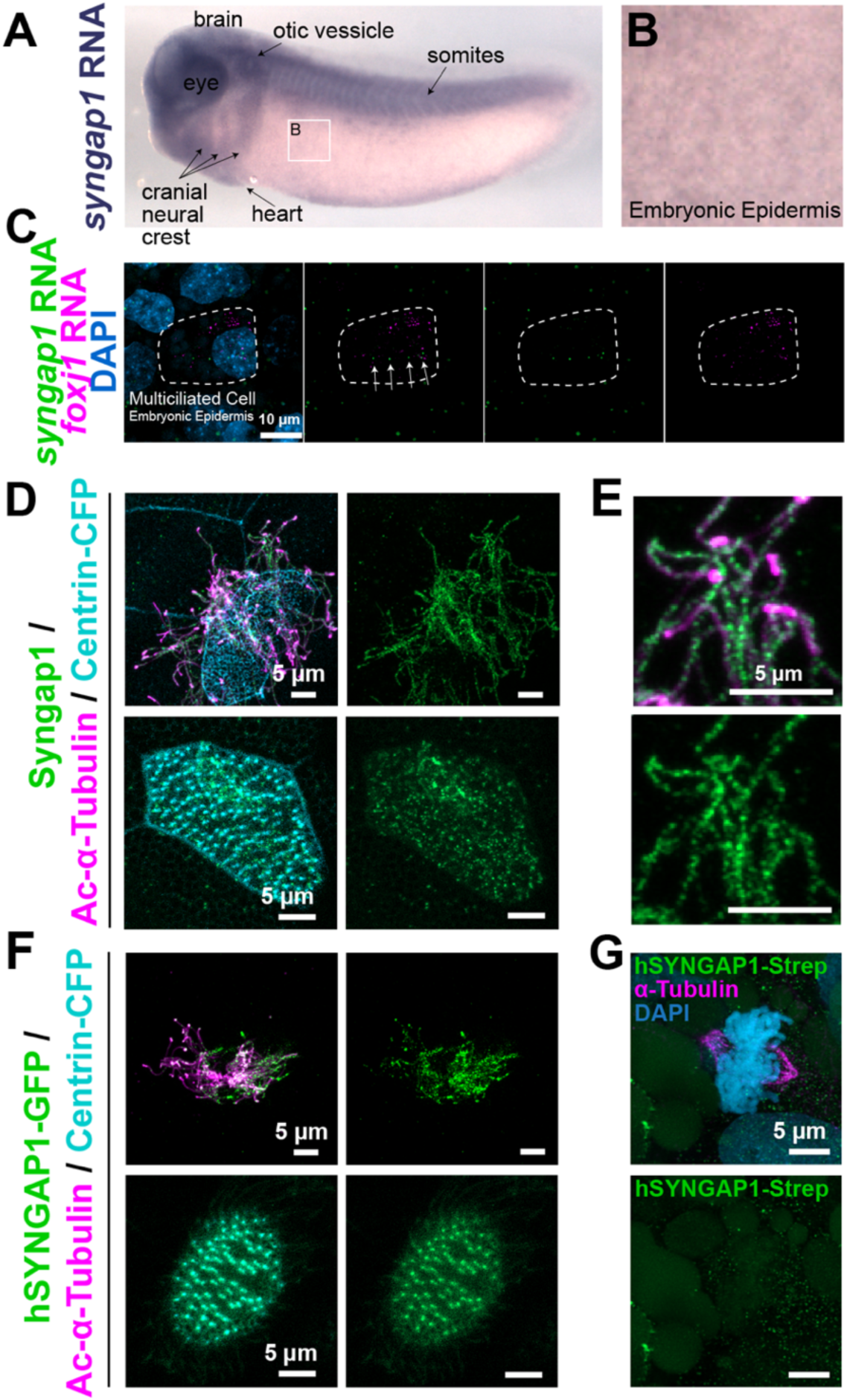
Syngap1 is highly expressed in *Xenopus* ciliated cells and localizes to cilia. (A) Whole mount RNA *in situ* hybridization for *syngap1* during *X. tropicalis* development. *syngap1* expression is enriched in the eye, brain, otic vesicle, and heart. (B) Higher magnification of the multiciliated embryonic epidermis, demonstrating *syngap1* expression. (C) HCR against *syngap1* and *foxj1* RNA in wildtype *Xenopus tropicalis* embryo multiciliated epidermis. *syngap1* RNA (green) is expressed in multiciliated cells expressing *foxj1* (magenta). Nucleus is labeled with DAPI. (D) Endogenous Syngap1 protein (antibody stain, green) localizes to the cilium (basal bodies labeled by centrin-CFP, cyan; axonemes labeled by acetylated alpha-Tubulin, magenta) in *Xenopus* multiciliated cells. (E) Higher magnification of D, showing axonemes in more detail. (F) GFP-tagged human SYNGAP1 (alpha-2 isoform, green) localizes to the cilium (basal bodies labeled by centrin-CFP, cyan; axonemes labeled by acetylated alpha-Tubulin, magenta) in *Xenopus* multiciliated cells. (G) Strep-tagged human SYNGAP1 (green) does not localize to spindle microtubules (labeled by alpha tubulin, magenta) in *Xenopus*. Nucleus is labeled with DAPI (blue).

**Figure S11.**
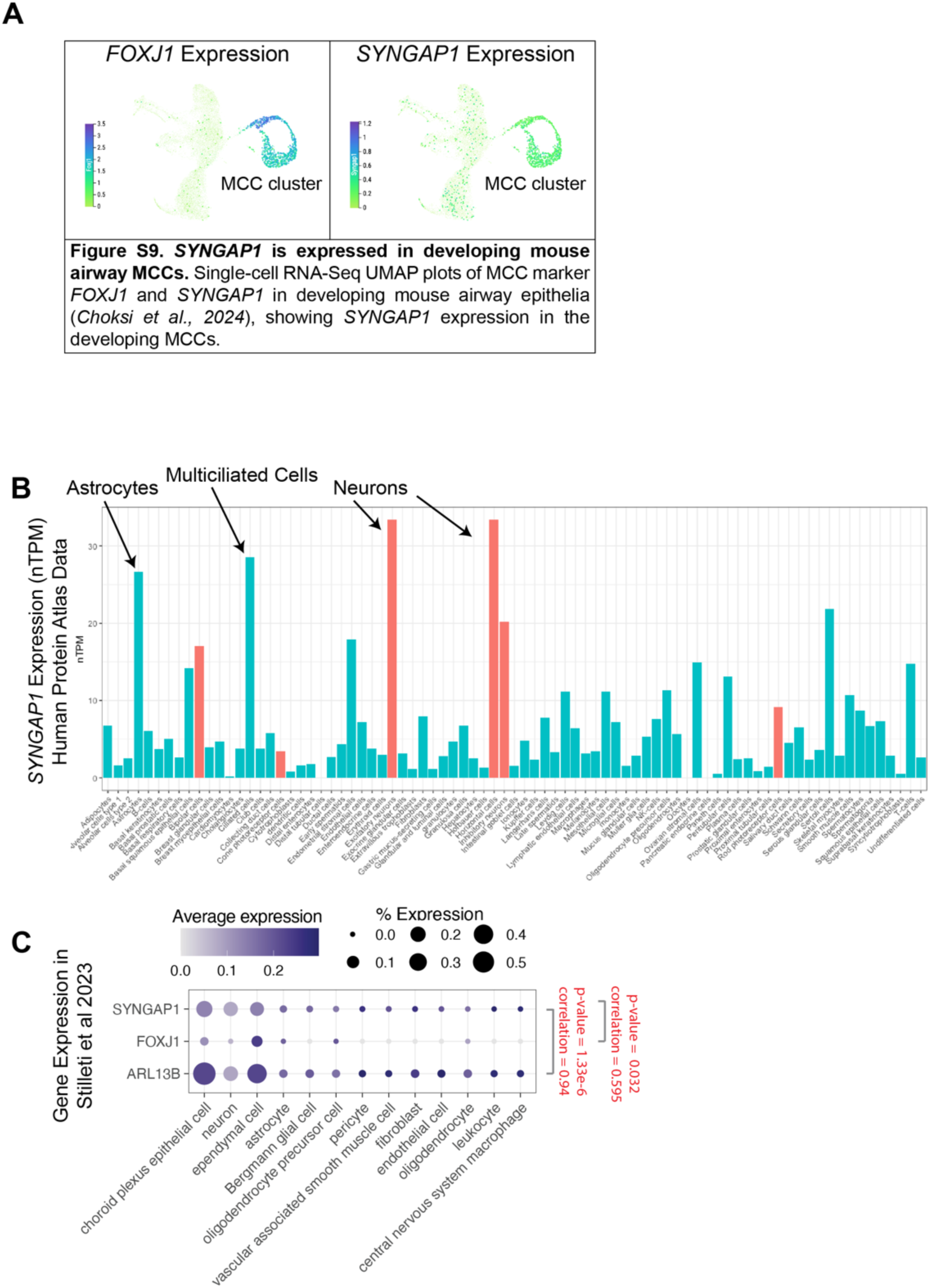
SYNGAP1 is expressed in developing mouse airway multiciliated cells (MCCs) and human ciliated cells. (A) Single-cell RNA-Seq UMAP plots of MCC marker *FOXJ1* and *SYNGAP1* in developing mouse airway epithelia (Choksi et al. 2024), showing *SYNGAP1* expression in the developing MCC lineage. (B) *SYNGAP1* normalized expression level across 81 cell types from Human Protein Atlas. Neuronal cell types are highlighted in red-orange and non-neuronal cell types are in turquoise. (C) *SYNGAP1* is significantly correlated with cilia markers *FOXJ1* and *ARL13B* in the *Siletti et al* dataset (Siletti et al. 2023).

**Figure S12.**
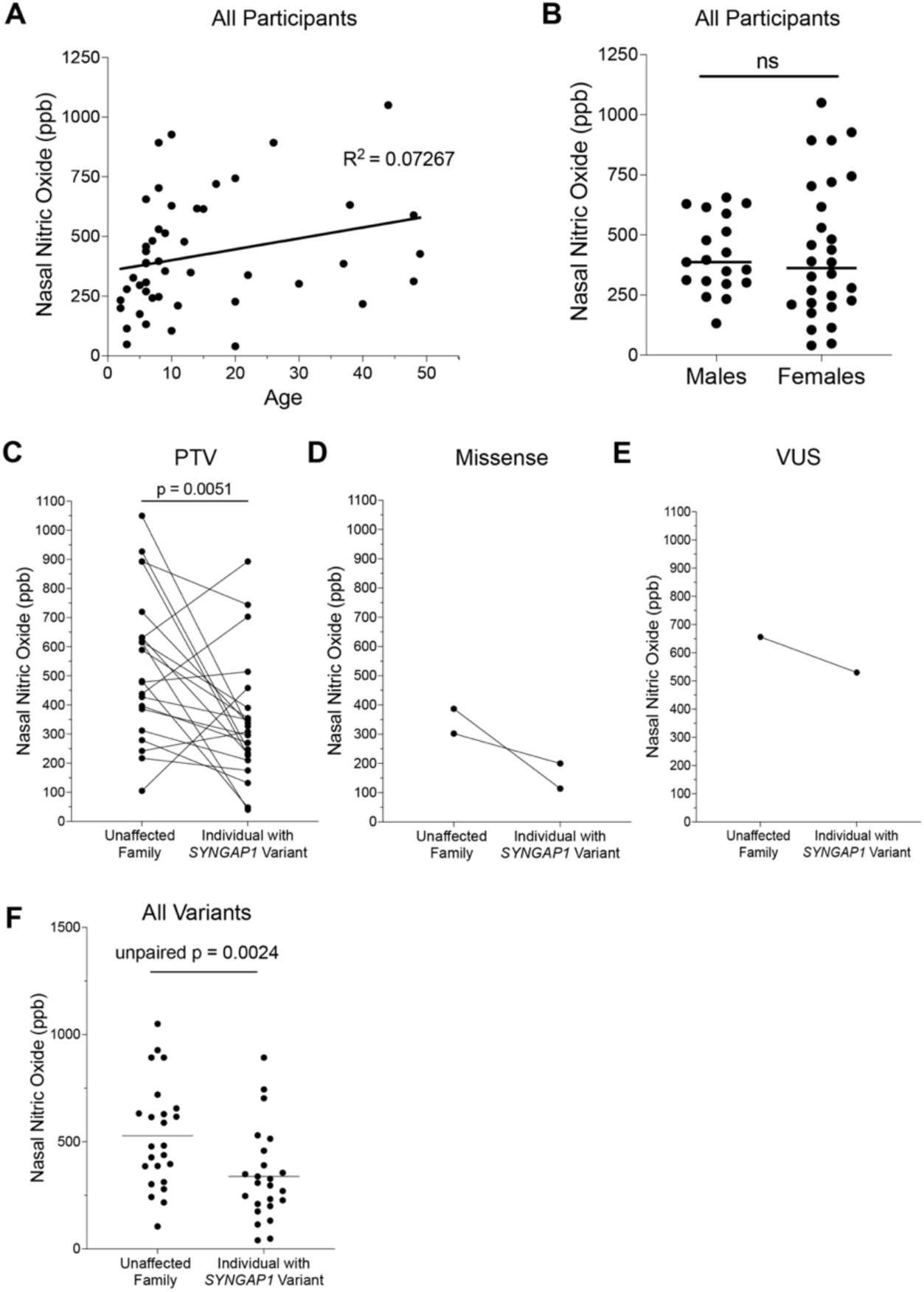
Sex and age differences do not explain the observed change in nasal nitric oxide levels in *SYNGAP1* patients. (A) There is no relationship between measured nasal nitric oxide levels nNO (NIOX VERO nNO machine) and age. (B) There is no relationship between nNO and sex assigned at birth. (C) Patients with protein truncating variants in *SYNGAP1* have significantly lower nasal nitric oxide levels (p = 0.0051, one-sided Wilcoxon matched-pairs signed rank test, NIOX VERO nNO machine) compared to immediate family members (sibling or parent). (D) Patients with missense variants in *SYNGAP1* have lower nasal nitric oxide levels compared to immediate family members (sibling or parent), n=2. (E) Patients with variants of unknown significance in *SYNGAP1* have lower nasal nitric oxide levels compared to immediate family members (sibling or parent), n=1. (F) Patients with *de novo* variants in *SYNGAP1* have significantly lower nasal nitric oxide levels on average when tested unpaired (p = 0.0024, one-sided Mann-Whitney rank sum test) compared to controls.

## Supplemental Table Legends

**Table S1. PPI Network Enrichment and Permutation Results.** Summary table with genesets, Fisher’s exact test results for enrichment of autism proteins in the four sets of cytoskeletal proteomes, and Fisher’s exact test results for permutation testing.

**Table S2. Expression correlations of autism genes with *ARL13B* or *FOXJ1* by cell type.** Cell types in the *Tabula Sapiens* dataset, their tissue of origin, and how many cells appear in that tissue; percent of each cell type that express a given gene for Tabula Sapiens 2023 and for Siletti 2023; tables with Pearson correlation of cilia markers (*FOXJ1*, *ARL13B*) for Tabula Sapiens data and tables with Pearson correlation of cilia markers (*FOXJ1*, *ARL13B*) for Siletti et al.

**Table S3. Summary of autism protein localization screen in primary cilia.** Summary table with results of the localization screen in motile and primary cilia.

**Table S4. Citizen patient medical record summary data.** Summary table of number and percentage of patients with non-benign variants with structural defects and heart murmurs; summary table of cardiac symptoms of patients with non-benign variants for each gene (separate tabs).

**Table S5. Summary of *SYNGAP1* morpholino and human SYNGAP1 rescue experiment.** Summary table with raw counts for the SYNGAP1 morpholino rescue experiment, breakdown of phenotype by percentages, Fisher’s exact test results, and pairwise Fisher’s exact test results.

